# Metadichol® induced expression of the TLR family of receptors in PBMCs

**DOI:** 10.1101/2024.04.04.588068

**Authors:** P.R. Raghavan

## Abstract

Toll-like receptors (TLRs) are fundamental components of the innate immune system and function as the primary sensors of invading pathogens by recognizing prevalent microbial compounds. The expression of all ten transcription factors belonging to the TLR family (1–10), with the exception of TLR4, exhibited an inverted U-shaped reaction to metadichol, a nanoemulsion composed of long-chain alcohol, as determined by the Q-RT□PCR assay. The response pattern suggested that low, moderate, and high concentrations directly affect TLR expression. This may indicate a dual-phase or dose-dependent effect on the immune system regarding TLR regulation. Compounds targeting TLRs can stimulate or inhibit these receptors, thereby affecting the immune response. Adjusting immunological activation, crucial for therapeutic purposes in conditions such as inflammation, cancer, infection, allergies, and autoimmune diseases, requires an inverse U-shaped response. Currently, no single-molecule examples can elicit the activation of every Toll-like receptor (TLR). This study examined the expression levels of all ten Toll-like receptors (TLRs), MYD88, and the downstream genes IRAK4, TRAF3, and TRIF. Metadichol expresses all the 15 genes.

## Introduction

The Toll receptor (TLR) family comprises ten receptors that are found either on the cell surface or in intracellular compartments (1). TLRs are key players in our immune system, and there is a need for them to maintain the normal functioning of our innate immune system. Targeting TLR signaling represents a new challenge for the treatment of many diseases. In the human body, TLR1, TLR2, TLR4, and TLR5 are intracellular compartments localized on the cell surface, whereas TLR3, TLR7, TLR8, and TLR9 are located in the endoplasmic reticulum, endosome, lysosome, or Endo lysosome compartments, respectively (1,2).

Selection has shaped the evolutionary lineage of the TLR gene family throughout vertebrates, mirroring the coevolutionary relationships between TLR proteins and their pathogenic counterparts (3). Evolutionary pressure has effectively preserved the integrity of TLRs, thereby facilitating their ability to identify and react to a wide range of pathogens. The number of Toll-like receptors (TLRs) in vertebrates varies considerably, from ten in humans to two hundred and twenty-two in purple sea urchins. This diversity underscores the evolutionary expansion and specialization of TLRs in the detection of a broad spectrum of microbial components [4].

### Structure of the TLR and Ligand Recognition

TLRs are distinguished by a transmembrane domain, a cytoplasmic Toll/IL-1 receptor (TIR) domain, and leucine-rich repeat (LRR) ectodomains that facilitate the recognition of pathogen-associated molecular patterns (PAMPs) [5]. Additionally, TLRs contain an LRR ectodomain that initiates subsequent signaling. The wide range of PAMPs that can be recognized by TLR structures, including lipids, lipoproteins, proteins, and nucleic acids, is crucial for the detection of various pathogens [6]. The adaptive evolution of TLRs is dependent on this structural versatility, which allows them to respond to a constantly shifting landscape of microbial threats.

### Mechanisms of Transcriptional Regulation and Signaling

Upon ligand binding, TLRs activate transcription factors such as interferon regulatory factors (IRFs) and NF-κB, which are essential for regulating the expression of genes implicated in antiviral immunity and inflammatory responses [7]. The recruitment of particular adaptor molecules is a component of signaling pathways, which demonstrates specificity and complexity. By maintaining a balanced immune response (8), this complex regulation prevents excessive inflammation, which may result in tissue injury.

### Disease-Related TLRs and Therapeutic Targeting

TLRs are known to have a substantial impact on the development of various diseases, such as malignancies, infectious diseases, and inflammatory conditions (9). Their participation in disease mechanisms has rendered them appealing therapeutic targets. An example of this is the correlation between susceptibility to diseases such as colorectal cancer and polymorphisms in TLR genes; this finding demonstrates the significance of genetic variations in relation to TLR function and disease risk (10). Therapeutic targeting of TLRs or their associated signaling pathways represents a potentially effective strategy for modulating immune responses in various diseases. Nonetheless, the complexity of the TLR signaling pathways and the specificity of ligand recognition present obstacles in the development of safe and efficacious therapeutics [11].

The TLR family of transcription factors serves as an intermediary between innate and adaptive immunity and is vital to the innate immune system (12). As a result of their evolutionary adaptation, vertebrates possess an advanced system for detecting and responding to pathogens. It is critical to comprehend the complex mechanisms underlying TLR signaling and their involvement in diseases to fully exploit their potential as therapeutic targets (13).

In addition to TLR family 1-10 and MYD88, the innate immune system’s response to pathogens is dependent on the Toll-like receptor (TLR) signaling pathway, which includes the genes IRAK4 (14-17), TRAF3 (18-20), TRIF (21-23), and TRAF6 (22-25). The collaboration of these genes guarantees an appropriate immune response against pathogens, underscoring the significance and intricacy of the TLR signaling pathway in the defense mechanisms of the host.

IRAK4: A crucial kinase, interleukin-1 receptor-associated kinase 4 (IRAK4), facilitates the transmission of signals from all TLRs, with the exception of TLR3. It is critical for the induction of subsequent signaling pathways that initiate the production of proinflammatory cytokines via mitogen-activated protein kinases (MAPKs) and nuclear factor kappa B (NF-κB).

TRAF3, which stands for tumor necrosis factor receptor-associated factor 3, has been identified as an antagonist of specific signaling pathways, such as the proinflammatory TLR and noncanonical NF-κB signaling pathways. Its degradation, which activates MyD88-dependent signaling and inhibits TRIF-dependent signaling, is essential for the regulation of both MyD88-dependent and TRIF-dependent signaling.

TRAF6 facilitates the transduction of signals from TLRs and other immune receptors as an E3 ubiquitin ligase. TRAF6 and IRAK form a complex in the MyD88-dependent TLR4 signaling pathway; this complex stimulates MAPKs and NF-κB, ultimately resulting in the synthesis of inflammatory cytokines.

TRIF: The adapter protein TIR-domain-containing adapter-inducing interferon-β (TRIF) is crucial for signaling pathways mediated by TLR3 and TLR4. It participates in the MyD88-independent pathway, which is responsible for inducing type I interferons (IFNs) and activating interferon regulatory factor 3 (IRF3). TRIF induces the activation of receptor-interacting serine/threonine kinase 1 (RIPK1) and TANK-binding kinase 1 (TBK1). This phosphorylation of IRF3 facilitates its translocation to the nucleus and subsequent synthesis of IFN1.

Research on the intricacies of TLR expression in PBMCs has shown that TLRs are expressed on various cell types within PBMCs, such as monocytes, dendritic cells, B cells, and T cells (26-28). TLR activation in PBMCs leads to the production of proinflammatory cytokines, activation of antigen presentation, and acquired immunity In autoimmune conditions such as systemic lupus erythematosus (SLE) and autoimmune thyroid disease (AITD), dysregulated TLR expression and activity have been observed, contributing to the pathogenesis of these diseases (29). Overall, TLRs play a crucial role in immune responses within PBMCs by detecting pathogens and modulating immune functions.

There are many small molecules that can express and activate Toll-like receptors (TLRs) in human cells

1. BCG: Acts on TLR2/4.
2. MPL: TLR4 Target.
3. CBLB502: Activates TLR5
4. Imiquimod: TLR7 stimulation.
5. IMO2055: Triggers TLR9.
6. MGN1703 Acts on TLR9.

These small molecules are designed to interact with specific TLRs, mimicking the action of natural ligands and inducing immune responses that can be beneficial in various therapeutic contexts, including vaccine development and immunotherapy (30).

The advantages of using small molecule Toll-Like Receptor (TLR) agonists over natural TLR ligands include the following:

1. Specificity: Small molecules can target specific TLRs, allowing for precise modulation of immune responses without affecting other pathways, unlike natural ligands that may activate multiple TLRs simultaneously.
2. Stability and Safety: Small molecule agonists are often more stable than natural ligands, reducing the risk of degradation and enhancing their safety profile for therapeutic applications.
3. Customization: Small molecules can be chemically modified to optimize their pharmacokinetic properties, bioavailability, and efficacy, providing flexibility in drug design and development.
4. Therapeutic potential: Small-molecule TLR agonists have shown promise in various applications, such as vaccine adjuvants, anticancer therapies, and immunotherapy, demonstrating their versatility and potential for clinical use. These advantages highlight the utility of small-molecule TLR agonists as valuable tools for modulating immune responses in a controlled and targeted manner for therapeutic purposes.

There is no single small molecule that activates or expresses the entire TLR family, including the MYD88 transcription factor family. Metadichol (31). treated PBMCs express all TLRs, and MYD88 is the first example of a small molecule displaying these properties.

### Experimental

All work was outsourced commercially to the service provider Skanda Life Sciences Pvt Ltd., Bangalore, India. The gene network analysis using Pathway Studio was outsourced commercially to ELSEVIER R&D Solutions, Inc. (32). The raw qRT□PCR data are provided in the Supplemental files.

### Cell line and cell conditions

#### 1. Isolation of Human WBCs

Preparation of the blood sample: Fresh human blood was collected in EDTA-containing tubes, diluted with PBS at a 1:1 ratio, and mixed by inverting the tube.

Isolation of Mononuclear Cells-In a 15 ml centrifuge tube, 5 ml of Histopaque-1077 was added, and 5 ml of the prepared blood was slowly layered on a histopaque from the edge of the tube without disturbing the histopaque layer. Then, the tubes were centrifuged at 400 × g for exactly 30 min at room temperature with brake-off settings. After centrifugation, the upper layer was discarded with a Pasteur pipette without disturbing the interphase layer. The interphase layer was carefully transferred to a clean centrifuge tube. The cells will be washed with 1X PBS and again centrifuged at 250 × g for 10 mins. (2X). After centrifugation, the supernatant was discarded, and the pellet was collected in RPMI media supplemented with 10% FBS (fetal bovine serum). Cells were counted, and viability was checked with a hemocytometer. After cell maintenance and seeding, media at a density of 1×106 cells/ml were prepared, and the cells were seeded into 6-well plates and incubated for 24 hr at 37°C with 5% CO2. After 24 h of seeding, the media was carefully removed, and the cells were treated with the indicated test samples (concentration was selected on the basis of the MTT experiment) and incubated for 24 h at 37°C in a CO2 incubator.

### Sample Preparation and RNA Isolation

The treated cells were dissociated, rinsed with sterile 1X PBS and centrifuged. The supernatant was decanted, and 0.1 ml of TRIzol was added and gently mixed by inversion for 1 min. Samples were allowed to stand for 10 min at room temperature. Then, 0.75 ml of chloroform was added to each 0.1 ml of TRIzol The contents were vortexed for 15 seconds. The tube was allowed to stand at room temperature for 5 mins. The resulting mixture was centrifuged at 12,000 rpm for 15 min at 4°C. The upper aqueous phase was collected in a new sterile microcentrifuge tube to which 0.25 ml of isopropanol was added, gently mixed by inverting the contents for 30 seconds and incubated at -20°C for 20 minutes. The contents were centrifuged at 12,000 rpm for 10 minutes at 4°C. The supernatant was discarded, and the RNA pellet was washed by adding 0.25 ml of 70% ethanol. The RNA mixture was centrifuged at 12,000 rpm at 4°C. The supernatant was carefully discarded, and the pellet was air-dried. The RNA pellet was then resuspended in 20 μl of DEPC-treated water. The total RNA yield (Table 2) was quantified using a Spectradrop (Spectramax i3x, Molecular Devices, USA).

**Table 1.** Treatment concentrations.

| Cell line | Sample name | Treatment details |
| --- | --- | --- |
| Human PBMC | Metadichol | Control |
|  |  | 1 pg |
|  |  | 100 pg |
|  |  | 1 ng |
|  |  | 100ng |

**Table 2.** RNA isolation yields.

| RNA yield (ng/μl) | Test concentrations | | | | |
| --- | --- | --- | --- | --- | --- |
|  | 0 | 1 pg/ ml | 100 pg/ ml | 1 ng/ ml | 100 ng/ ml |
| Human PBMC's | 438.00 | 304.64 | 161.92 | 376.24 | 382.88 |

### Q-RT □ PCR analysis cDNA synthesis

cDNA was synthesized from 500 ng of RNA using a cDNA synthesis kit from the Prime Script RT Reagent Kit (TAKARA) with oligo dT primers according to the manufacturer s instructions. The reaction volume was set to 20 μl, and cDNA synthesis was performed at 50°C for 30 min, followed by RT inactivation at 85°C for 5 min using the applied biosystem Veritii. The cDNA was further used for real-time PCR analysis.

### Primers and qPCR analysis

The PCR mixture (final volume of 20 μl) contained 1.4 μl of cDNA, 10 μl of SyBr Green Master Mix and 1 μl of the respective complementary forward and reverse primers (Table 3) specific for the respective target genes. The reaction was carried out with enzyme activation at 95°C for 2 minutes, followed by a 2-step reaction with an initial denaturation and annealing step at 95°C for 5 seconds, annealing for 30 seconds at the appropriate temperature, and amplification for 39 cycles followed by secondary denaturation at 95°C for 5 seconds, with 1 cycle of melt curve capture ranging from 65°C to 95°C for 5 sec each. The obtained results were analyzed, and the fold changes in expression or regulation were calculated.

**Table 3.**
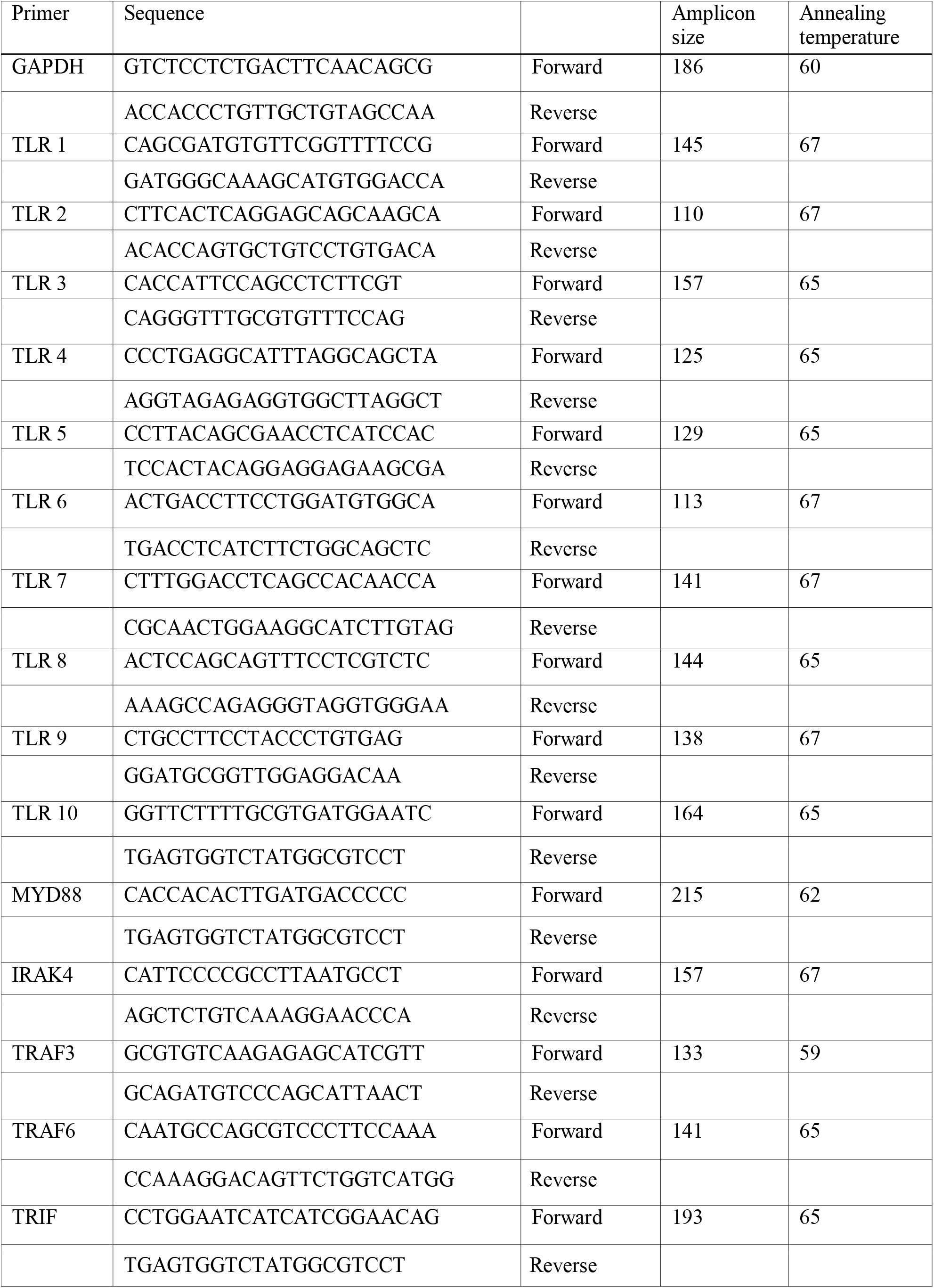
Primer Details.

## Results and Discussion

Our results revealed that the expression of TLR1-10 and the expression of MYD88 were associated with an inverted U-shaped response, with the exception of TL4, whose expression was downregulated (Figure 1). The inverted U-shaped gene expression response to varying concentrations of a small molecule can be attributed to complex regulatory mechanisms. This response pattern, also known as a biphasic dose response, suggests that low and high concentrations of the small molecule may affect gene expression differently. The literature (33) provides insights into the origin and dynamics of inverted U/U dose-response relationships, which emerge due to interactions among various factors. Some examples (34) that exhibit an inverted U-shaped response in expression include genes regulated by estrogen (E2). Research has shown that the effects of estrogen on hippocampal activity in young women can exhibit both a monotonically increasing relationship and an inverted U-shaped dose-response function. The response of genes in the hippocampus to estrogen levels can follow an inverted U-shaped pattern, indicating a complex relationship between estrogen signaling and gene expression.

**Figure 1.**
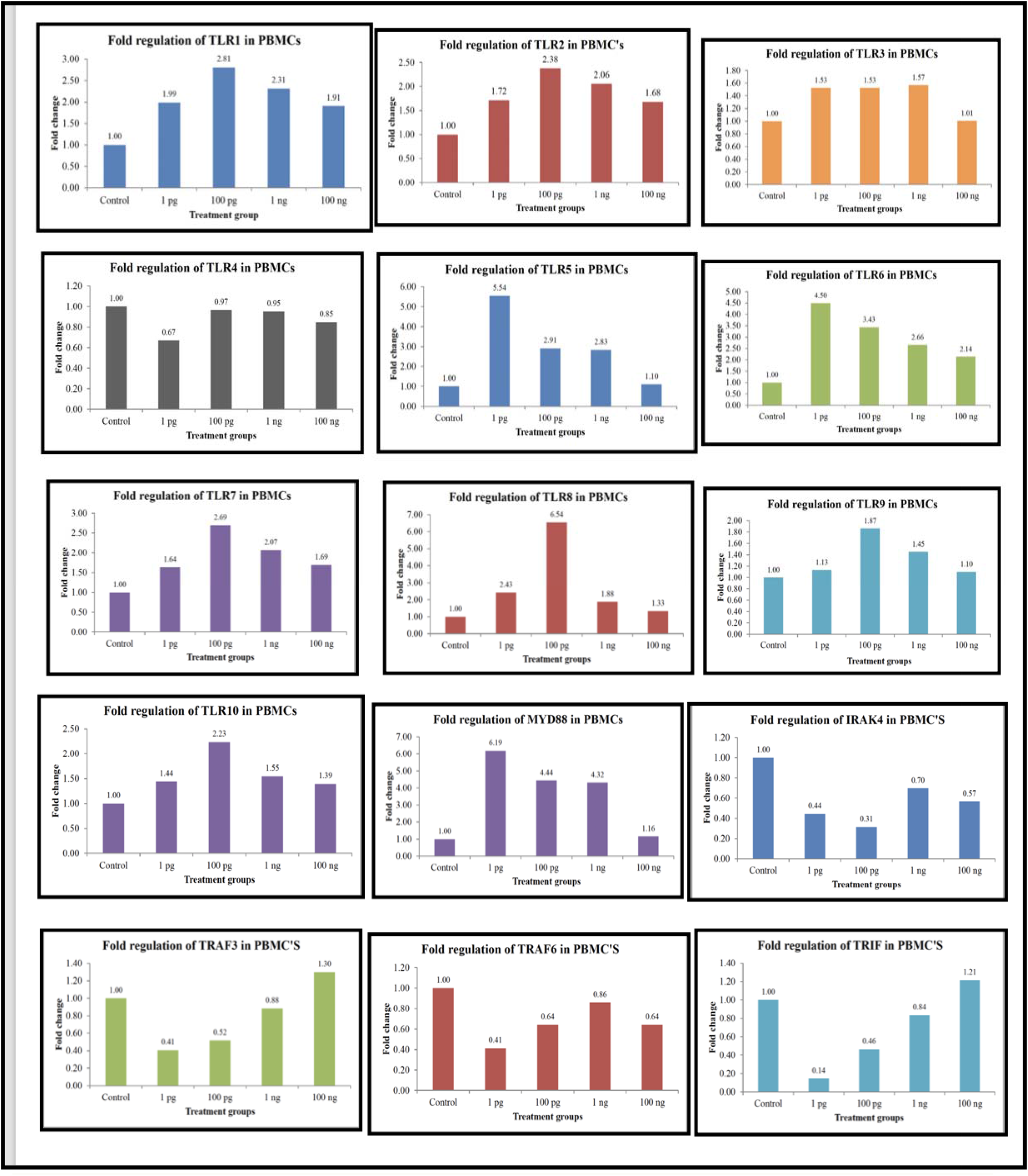
Q-RT□PCR Fold Changes.

Studies have suggested that the dosage of antibiotics can affect the inverted U-shaped response in bacterial resistance genetics (35). Gene expression related to antibiotic resistance mechanisms may exhibit a nonlinear response pattern. These examples highlight the diverse nature of genes that can display an inverted U-shaped response in expression, reflecting the intricate regulatory mechanisms governing gene expression in response to varying concentrations of small molecules.

There are limited examples of drugs that exhibit an inverted U-shaped dose□response function and have been approved for therapeutic use. However, one example is fenfluramine, a 5-HT releaser, which is used to treat seizures associated with Lennox–Gastaut syndrome and Dravet syndrome (36). This example highlights how certain drugs with inverted U-shaped dose-response curves can have therapeutic applications, emphasizing the importance of understanding dose-response relationships in clinical settings. While specific small molecules can target individual TLRs, the concept of a single small molecule affecting all TLR receptors simultaneously is complex due to the structural and functional diversity among TLRs. We have previously shown that in fibroblasts treated with metadichol, all 49 nuclear receptors are expressed (37).

Based on the known functions of nuclear receptors and TLRs, it is more likely that the activation of nuclear receptors may indirectly modulate TLR expression rather than directly increase it. Nuclear receptors regulate various immune responses, including inflammation, and can interact with transcription factors that control TLR-ind uced gene expression (38,39).

Liver X receptors (LXRs), including LXRα and LXRβ, play crucial roles in regulating Toll-Like Receptor (TLR) signaling pathways and inflammation through various mechanisms (40). Regulation of TLR Signaling: LXRs inhibit signaling from TLRs 2, 4, and 9 to downstream NF-κB and MAPK effectors through ABCA1-dependent changes in membrane lipid composition.Gene expression control: LXRs control the expression of genes involved in cholesterol, fatty acid, and phospholipid metabolism by directly binding to LXR response elements (LXREs) in target promoters. They can activate lipid metabolism-related genes and antagonize inflammatory gene expression triggered by TLR activation Anti-Inflammatory Effects: Activation of LXRs leads to the repression of inflammatory gene expression both in cells and in mice. This anti-inflammatory effect is primarily mediated through the regulation of the Abca1 sterol transporter, altering membrane cholesterol homeostasis and inhibiting the NF-κB and MAPK signaling pathways downstream of TLRs (41). In summary, LXRs, particularly LXRα and LXRβ, link metabolism to inflammation by regulating membrane composition, TLR signaling, and inflammatory gene expression, highlighting their role in modulating immune responses.

The nuclear receptor aryl hydrocarbon receptor (AhR) plays a crucial role in modulating Toll-like receptor (TLR)-induced gene expression. Activation of AhR can lead to the downregulation of microRNAs targeting anti-inflammatory and myeloid-derived suppressor cell (MDSC)-regulatory genes, resulting in the induction of chemokines and their receptors. This process ultimately leads to upregulating proinflammatory responses in inflammatory diseases (42). Studies have shown that AhR activation can attenuate inflammation through various mechanisms, such as thymic atrophy, apoptosis, regulatory T-cell (Treg) induction, myeloid-derived suppressor cell (MDSC) modulation, cytokine suppression, and epigenetic changes. By regulating these processes, AhR activation can protect against inflammatory diseases such as colitis, multiple sclerosis, atopic dermatitis, and psoriasis (43). Furthermore, AhR activation has been implicated in regulating the differentiation of inflammatory dendritic cells, which may contribute to the immune response under inflammatory conditions. Metadichol is a inverse agonist of the AHR nuclear receptor (44)

Other receptors, such as the PPAR and GR (glucocorticoid receptor), also regulate TLRs (45). GRs are involved in regulating the immune response and can interact with TLR signaling pathways. They modulate inflammation and immune cell function in response to TLR activation.

PPARs are another group of nuclear receptors that interact with TLR signaling pathways. They can negatively regulate TLR-induced inflammation by inhibiting the expression of proinflammatory cytokines and promoting anti-inflammatory responses. Negative regulation of TLR signaling also occurs, via the orphan nuclear receptor SHP (short heterodimer partner also known as NROB2) has been shown to act as a negative regulator in inflammatory signaling triggered by TLRs (23)

Retinoic acid receptor-related orphan receptors (RORs): RORs are nuclear receptors that have been implicated in the regulation of TLR signaling. They can modulate the expression of genes involved in the immune response, including some TLR genes (46).

The crosstalk between nuclear receptors and TLR signaling pathways can influence the expression of genes involved in immune responses, but the direct impact of nuclear receptor activation on TLR expression may involve complex regulatory mechanisms that require further investigation.

One exception to this inverted U-shaped response was TLR4, which was not comparable to the downregulated controls. This inhibition of TLR4 (47,48) has been associated with the potential treatment of various diseases. Research suggests that inhibiting TLR4 signaling pathways can attenuate inflammatory responses and cardiac myocyte apoptosis. It is a potential target for treating conditions such as trauma-hemorrhage, sepsis, acute lung injury, chronic inflammatory arthritis, airway inflammation, and autoimmune diseases.

Additionally, TLR4 inhibition has been studied in the context of myocardial inflammation, cardiovascular disease, allergic diseases, obesity-associated metabolic diseases, and neuronal degeneration. TLR4 has also been reported to be associated with age-related diseases, further highlighting its potential significance in disease research (49). Furthermore, TLR4 inhibitors are being researched for their potential in the treatment of severe COVID-19 and related disorders (50.51). Interestingly, metadichol is a powerful inhibitor of the SARS-CoV-2 virus (52). The inhibition of TLR4 has been studied under various conditions, including insulin resistance and inflammatory diseases.

Research has shown that TLR4 inhibition can have protective effects against lipid-induced insulin resistance in skeletal muscle. For instance, the selective TLR4 inhibitor TAK-242 has been found to protect against lipid- and LPS-induced insulin resistance in muscle cells and rats (53).

TLR4 inhibition has been linked to protection against sensory and motor dysfunction in an animal model of autoimmune peripheral neuropathy (39). Furthermore, in the context of Alzheimer’s disease, inhibiting TLR4 has been associated with neuroprotection and the modulation of microglial polarization (54).

TLR4 can promote cancer growth and chemoresistance in epithelial ovarian cancer and is correlated with metastatic potential in prostate cancer. In breast cancer, TLR4 has been associated with increased migration, invasion, angiogenesis, and metastatic potential of cancer cells. While TLR4 activation on immune cells can promote antitumor responses, its activation on tumor or stromal cells may enhance tumor progression (55,56). Eritoran, a synthetic analog of lipid A from Rhodobacter sphaeroides, is known to inhibit TLR4 by preventing the interaction between TLR4 and lipid A. Bacterial LPS-induced colon cancer can be prevented by the administration of Eritoran (57).

In addition, TLR agonists play important roles in the activation of both the innate and adaptive immune systems. TLR agonism has been shown to reduce tumor growth in treatment groups receiving combinations of therapeutic agents (58,59). The Calmette–Guerin strain (BCG) activates TLR2, TLR4, and TLR9. This activation of TLRs in urothelial cell carcinomas reduces cell death and decreases proliferation as well as metastasis. (60)

The expression of TLR5 on cancer cells decreases cell growth in breast cancer (61,62). Activation of TLR9 has an inhibitory effect on human glioma cell lines [47]. Such effects can sensitize cancerous cells to radiation treatment (63). The use of TLR agonists approved by the FDA (Food and Drug Administration, USA) for cancer treatment include BCG (which activates TLR2, TLR3, TLR4, and TLR9) and imiquimod (a TLR7 agonist) (64). TLRs play a role in mitigating inflammation, cell proliferation, apoptosis and chemoresistance in cancerous cells (65,66).

Another TLR of interest is TLR10, which is a unique Toll-like receptor known for its anti-inflammatory properties. Unlike other TLRs, TLR10 has been shown to act as an inhibitory receptor with suppressive effects (67). TLR10 can suppress inflammatory signaling in primary human cells, leading to a reduction in the production of inflammatory cytokines. This anti-inflammatory role of TLR10 makes it a potential target for therapeutic interventions aimed at modulating immune responses in conditions where excessive inflammation is detrimental. Despite the challenges posed by conflicting research findings regarding its function, further exploration of TLR10 mechanisms and signaling pathways could unveil new opportunities for developing targeted therapies that harness its unique anti-inflammatory properties (68).

TLR10 exhibits anti-inflammatory properties, activating gene transcription through MyD88. They act as coreceptors with other TLRs and share ligands with each other (55). Additionally, a functional variant of TLR10 has been associated with modifying NF-κB activity, impacting disease severity in conditions such as rheumatoid arthritis (69-72).

TLR1 surface expression decreases by 36% in older adults compared to young adults, indicating an age-associated defect in TLR1 function (73-74). TLR1 plays a role in aging by being associated with a decrease in surface expression on monocytes of older adults, which correlates with decreased cytokine production. Studies have shown that human TLR1/2 function is substantially diminished in the context of aging, with TLR1 surface expression being significantly lower in older adults than in young adults. This age-associated defect in TLR1 function may contribute to impaired immune responses to infections and age-related changes in immune signaling pathways. Increased TLR1 expression offers hope for reversing aging processes.

The highest fold change in expression was that of TLR8, which was greater than 6.5-fold. This increased expression of TLR8 in peripheral blood mononuclear cells (PBMCs) can have significant immunological implications (75). These proteins play a crucial role in the innate immune system by recognizing pathogen-associated molecular patterns (PAMPs) and initiating immune responses.

The following are some potential implications of increased TLR8 expression in PBMCs (76-78).

1. Enhanced Immune Response: TLR8 activation can lead to the production of various cytokines and chemokines, enhancing the immune response against pathogens
2. Autoimmune Diseases: Abnormal activation or overexpression of TLR8 may contribute to the pathogenesis of autoimmune diseases by recognizing self-RNA and promoting inflammation.
3. Cancer Immunotherapy: TLR8 is being explored as a target for cancer immunotherapy. Agonists of TLR8 can potentially enhances antitumor immunity by activating immune cells
4. Vaccine Adjuvants: TLR8 agonists are considered potential vaccine adjuvants for improving vaccine efficacy by stimulating a stronger immune response
5. Disease Biomarkers: The expression levels of TLR8 could serve as biomarkers for certain diseases, including infections and autoimmune disorders. Increased TLR8 expression in PBMCs is a marker of immune activation and could be leveraged for therapeutic purposes.

We further analyzed gene expression using one network analysis. The 15 genes were imported into Pathway Studio software (79,80) to perform gene set enrichment analysis (GSEA) against the Elsevier proprietary pathway collection GSEA, which revealed that the gene list was enriched with genes. These genes have more interactions among themselves than expected for a gene set of the same size and distribution degree randomly selected from the genome. This enrichment indicates that this set of genes shares a significant biological connection (81).

Figure 2 shows all relationships between 15 genes available in the Elsevier Biology Knowledge Graph. Analysis of gene networks using Pathway Studio and protein–protein interaction maps indicated the formation of a loop feedback network, as shown in Fig. (2). TLR’s regulate each and interact with VDR as we see in Figure 3.

**Figure 2.**
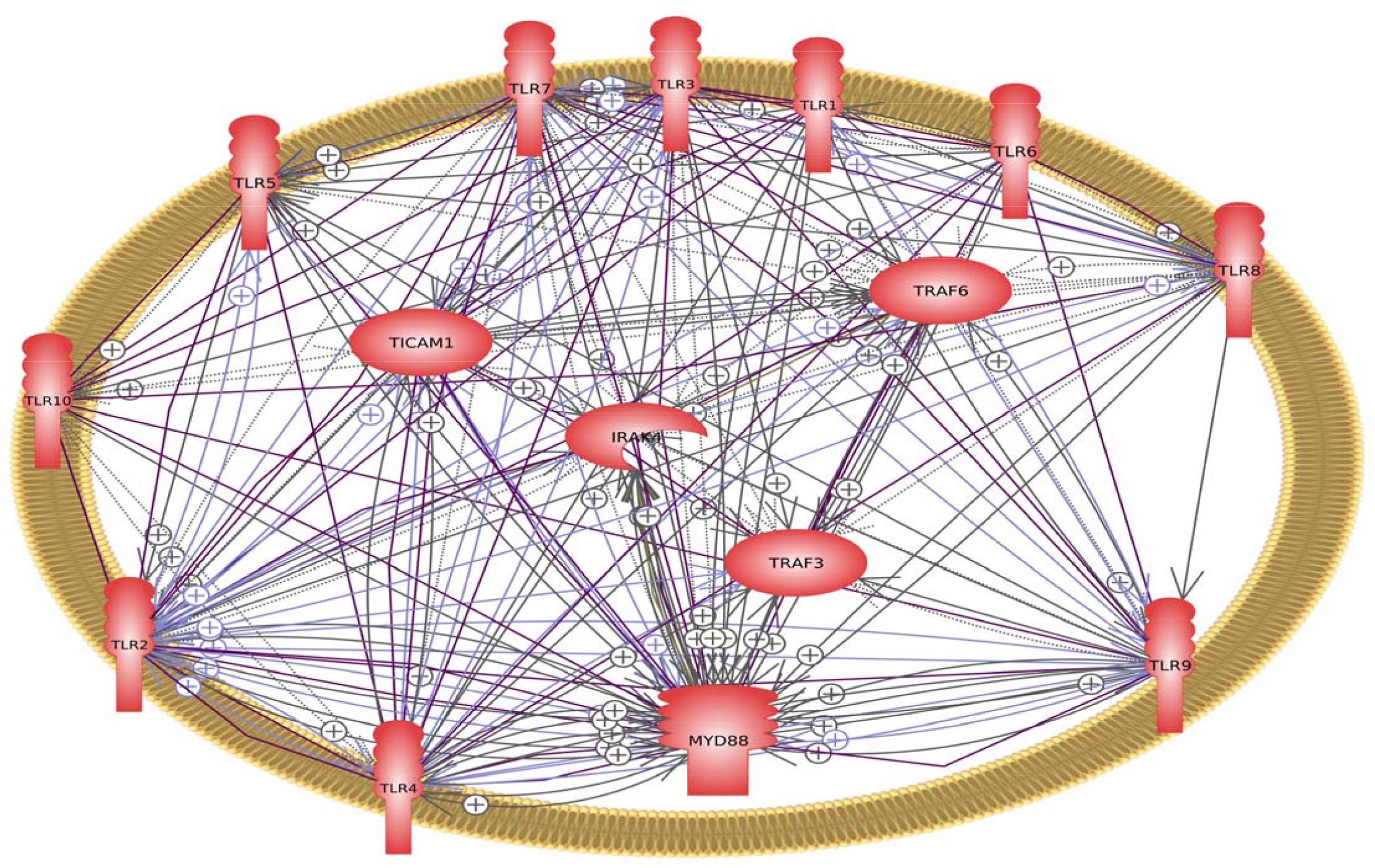
The interaction and regulatory network gene sets TLR1-10, MYD88, IRAK4, TRAF6, TRAF3, and TICAM1 (TRIF)

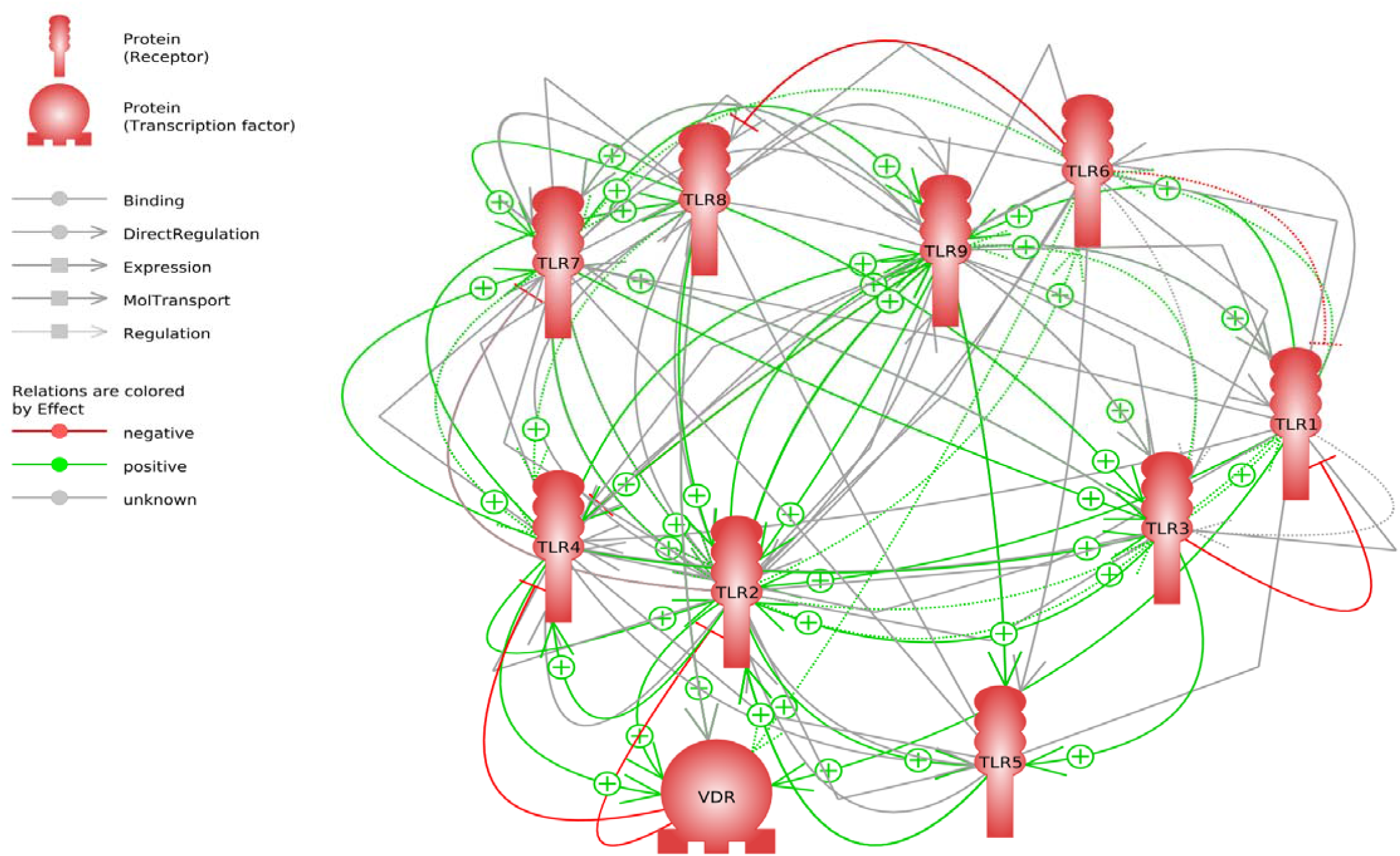

The TLR family of genes forms a closed loop network with the VDr. We see here that VDR down regulates TLR 4 which we have show with Q-RT-PCR. TLR6 regulates TLR8, TLR 3 down regulates TLR1 and VDR regulates TLR2 and TLR2 regulates TLR 7 (follow the red lines)

The TLR family of genes activates and regulates each other through intricate signaling pathways involving specific adaptor molecules. TLRs signal by recruiting adaptors such as MyD88, TRIF, TIRAP/MAL, or TRAM, leading to the activation of the transcription factors NF-κB and IRFs, which control the immune response (1,6,13).

**Tables 4-9** show key pathways and cell process diseases and organs that involve the TLR family of transcription factors.

**Table 4.** Biological process.

| Biological Process | Parent Folder | # of Entities | Expanded # of Entities | Overlap | Percent Overlap | Overlapping Entities | p-value |
| --- | --- | --- | --- | --- | --- | --- | --- |
| Antiviral Signaling through Pattern Recognition Receptors | Antigen Recognition in Innate Immune System; Antigen Recognition in Innate Immune System (Immunological Pathways); COVID19 induced immune response; generic; generic; plasma membrane; secretory vesicle | 31 | 36 | 10 | 27 | TRAF3, TRAF6, TLR2, TLR7, TICAM1, TLR3, TLR4, TLR8, MYD88, IRAK4 | 5.45964E-23 |
| Infection->TLRs in corneal ulcer | Cornea epithelium-immune sytem interactions | 95 | 281 | 13 | 4 | TLR7, TICAM1, TLR8, TRAF3, TRAF6, TLR6, IRAK4, TLR9, MYD88, TLR1, TLR2, TLR3, TLR4 | 6.35177E-20 |
| Fatty Acids Block TLR Signaling in Enteroendocrine Cell | Eating Behavior and Metabolism Regulation; Eating Behavior and Metabolism Regulation (Biological Process); digestive system; epithelium | 21 | 69 | 9 | 13 | TLR7, TLR9, TLR8, TLR10, MYD88, TRAF6, TLR3, TLR5, IRAK4 | 2.34166E-17 |
| Toll-like Receptors Act through MYD88 Signaling | Antigen Recognition in Innate Immune System; Antigen Recognition in Innate Immune System (Immunological Pathways); generic; generic; plasma membrane; secretory vesicle | 28 | 39 | 8 | 20 | TRAF6, TLR7, TLR5, TLR9, TLR8, TLR10, MYD88, IRAK4 | 4.31449E-17 |
| Toll-like Receptors in Sterile Inflammation | Inflammation Initiation; Inflammation Initiation (Inflammation Pathways); generic; generic | 75 | 103 | 9 | 8 | TICAM1, TRAF3, TRAF6, IRAK4, TLR9, MYD88, TLR2, TLR3, TLR4 | 1.02082E-15 |
| Mast-Cell Activation without Degranulation through IL1R1 and TLR Signaling | Mast Cells Activation; Mast Cells Activation (Myeloid Cells Activation in Inflammation); generic; hematological system | 27 | 155 | 9 | 5 | TLR7, TLR8, TRAF6, IRAK4, TLR9, TLR10, MYD88, TLR3, TLR5 | 4.45974E-14 |
| AXL Receptor Inhibits Macrophages and Dendritic Cells Function | Monocytes Activation; Monocytes Activation (Myeloid Cells Activation in Inflammation); generic; hematological system | 46 | 62 | 7 | 11 | TICAM1, TLR9, MYD88, TRAF6, TLR3, TLR4, IRAK4 | 4.98267E-13 |
| Mast-Cell Activation without Degranulation | Mast Cells Activation; Mast Cells Activation (Myeloid Cells Activation in Inflammation); generic; hematological system | 63 | 214 | 9 | 4 | TLR7, TLR8, TRAF6, IRAK4, TLR9, TLR10, MYD88, TLR3, TLR5 | 8.44955E-13 |
| Macrophage M1 Lineage | Monocytes Activation; Monocytes Activation (Myeloid Cells Activation in Inflammation); generic; hematological system | 66 | 345 | 10 | 2 | TLR7, TICAM1, TLR8, TRAF6, IRAK4, TLR9, TLR10, MYD88, TLR3, TLR5 | 1.09802E-12 |
| I-cell: CCK-8 Secretion and Regulation of Eating Behavior | Cells of Gastrointestinal Tract Activation; Cells of Gastrointestinal Tract Activation (Eating Behavior and Metabolism Regulation); digestive system; epithelium | 54 | 231 | 7 | 3 | TRAF6, IRAK4, TLR9, MYD88, TLR2, TLR4, TLR5 | 5.81468E-09 |

**Table 4;**The key biological pathways are shown in Table 4, and the most enriched pathway was signaling through pattern recognition receptors as well as through the MYD88 signaling pathway.

**Table 5;** shows cell processes that include antigen recognition, the immune response and cytokine pathways, among others.

**Table 5.** Key signaling pathways.

| pathways | # of Entities | Expanded # of Entities | Overlap | Percent Overlap | Overlapping Entities | p-value |
| --- | --- | --- | --- | --- | --- | --- |
| Antiviral Signaling through Pattern Recognition Receptors | 31 | 36 | 10 | 27 | TRAF3, TRAF6, TLR2, TLR7, TICAM1, TLR3, TLR4, TLR8, MYD88, IRAK4 | 5.20380E-25 |
| Toll-like Receptors (TLR) in Antiviral Innate Immune Response | 30 | 48 | 10 | 20 | TLR7, TICAM1, TLR9, TLR8, MYD88, TRAF6, TLR2, TLR3, TLR4, IRAK4 | 1.33514E-23 |
| COVID19+>AGER feed forward loop in innate immunity | 30 | 38 | 9 | 23 | TRAF6, TLR6, TLR2, TLR7, TLR4, TLR9, TLR8, MYD88, IRAK4 | 1.03891E-21 |
| Dendritic Cell Activation in Systemic Lupus Erythematosus | 73 | 242 | 12 | 4 | TLR7, TICAM1, TLR8, TRAF3, TRAF6, TLR6, IRAK4, TLR9, MYD88, TLR2, TLR3, TLR4 | 7.25697E-21 |
| Dendritic Cell Activation in COVID19 | 76 | 247 | 12 | 4 | TLR7, TICAM1, TLR8, TRAF3, TRAF6, TLR6, IRAK4, TLR9, MYD88, TLR2, TLR3, TLR4 | 9.32148E-21 |
| Plasmacytoid Dendritic Cell in Diabetes Mellitus Type 1 | 53 | 91 | 9 | 9 | TLR7, TICAM1, TRAF6, IRAK4, TLR9, MYD88, TLR2, TLR3, TLR4 | 4.91915E-18 |
| Apoptotic Keratinocytes Clearance Recession in Systemic Lupus Erythematosus | 79 | 379 | 11 | 2 | TICAM1, TRAF6, TLR6, IRAK4, TLR9, TLR10, MYD88, TLR1, TLR2, TLR4, TLR5 | 2.56647E-16 |
| Toll-like Receptors (TLR) in Periodontitis | 39 | 78 | 8 | 10 | TLR6, TICAM1, MYD88, TRAF6, TLR1, TLR2, TLR4, IRAK4 | 3.93659E-16 |
| Dendritic Cells Function in Atherosclerosis | 53 | 80 | 8 | 10 | TLR6, TLR7, TICAM1, TLR9, MYD88, TRAF6, TLR4, IRAK4 | 4.86359E-16 |
| IL36 Receptor Antagonist (IL1F5) Deficiency (DITRA) | 37 | 165 | 9 | 5 | TLR7, TLR8, TRAF6, IRAK4, TLR9, TLR10, MYD88, TLR3, TLR5 | 1.22898E-15 |

**Table 6**: shows protein regulation of cell processes..

**Table 6.** Protein regulators of Cell processes.

| Protein regulators of | Total # of Neighbors | Overlap | Percent Overlap | Overlapping Entities | p-value |
| --- | --- | --- | --- | --- | --- |
| apoptosis | 13743 | 15 | 0 | TLR3, TLR4, TLR10, TLR5, MYD88, IRAK4, TLR9, TRAF3, TRAF6, TLR1, TLR8, TLR6, TLR2, TLR7, TICAM1 | 0.00000E+00 |
| cell proliferation | 15770 | 14 | 0 | TLR3, TLR4, TLR5, MYD88, IRAK4, TLR9, TRAF3, TRAF6, TLR1, TLR8, TLR6, TLR2, TLR7, TICAM1 | 0.00000E+00 |
| PB-cell response | 599 | 13 | 2 | TLR3, TLR4, TLR10, TLR5, MYD88, TLR9, TRAF3, TLR1, TLR8, TLR6, TLR2, TLR7, TICAM1 | 2.62383E-16 |
| immunostimulation | 1326 | 15 | 1 | TLR3, TLR4, TLR10, TLR5, MYD88, IRAK4, TLR9, TRAF3, TRAF6, TLR1, TLR8, TLR6, TLR2, TLR7, TICAM1 | 8.70939E-16 |
| myeloid dendritic cell activation | 35 | 7 | 19 | TLR3, TLR4, TLR9, MYD88, TLR8, TLR2, TLR7 | 2.89483E-15 |
| intracellular signaling cascade | 1027 | 14 | 1 | TLR3, TLR4, TLR5, MYD88, IRAK4, TLR9, TRAF3, TRAF6, TLR1, TLR8, TLR6, TLR2, TLR7, TICAM1 | 3.37116E-15 |
| macrophage response | 735 | 13 | 1 | TLR3, TLR4, TLR10, TLR5, MYD88, TLR9, TRAF3, TRAF6, TLR1, TLR6, TLR2, TLR7, TICAM1 | 3.76778E-15 |
| immune response-regulating signaling pathway | 753 | 13 | 1 | TLR3, TLR4, TLR5, MYD88, IRAK4, TLR9, TRAF3, TRAF6, TLR8, TLR6, TLR2, TLR7, TICAM1 | 5.16000E-15 |
| erythrocyte apoptosis | 45 | 7 | 15 | TLR3, TLR4, TLR5, MYD88, TRAF6, TLR2, TLR7 | 1.94327E-14 |
| response to viruses | 1179 | 14 | 1 | TLR3, TLR4, TLR10, TLR5, MYD88, IRAK4, TLR9, TRAF3, TRAF6, TLR8, TLR6, TLR2, TLR7, TICAM1 | 2.32795E-14 |

**Table 7**: TLR’s presence in organs, including the testis, cerebral cortex, brain, neurons, lung, kidney, and liver, and these have diverse implications for immune responses and disease development.

**Table 7.** Diseases regulation by the gene set; TLR1-10, MYD88, IRAK4, TRAF6, TRAF3, and tICAM1 (TRIF)

| Protein regulators of | Total # of Neighbors | Overlap | Percent Overlap | Overlapping Entities | p-value |
| --- | --- | --- | --- | --- | --- |
| genital herpes | 35 | 11 | 30 | TLR3, TLR4, TLR9, TLR10, TLR5, MYD88, TLR1, TLR8, TLR6, TLR2, TLR7 | 1.28731E-29 |
| tibial dyschondroplasia | 38 | 7 | 17 | TLR3, TLR4, TLR5, MYD88, TRAF6, TLR2, TLR7 | 8.05934E-17 |
| liver disease | 1053 | 13 | 1 | TLR3, TLR4, TLR5, MYD88, IRAK4, TLR9, TRAF3, TLR1, TLR8, TLR6, TLR2, TLR7, TICAM1 | 1.69120E-16 |
| viral myocarditis | 186 | 9 | 4 | TLR3, TLR4, TLR9, TLR5, MYD88, IRAK4, TLR6, TLR2, TLR7 | 3.53879E-16 |
| myocarditis | 552 | 11 | 1 | TLR3, TLR4, TLR9, TLR5, MYD88, IRAK4, TLR1, TLR6, TLR2, TLR7, TICAM1 | 9.36950E-16 |
| immunopathology | 942 | 12 | 1 | TLR3, TLR4, TLR10, TLR5, MYD88, TLR9, TRAF3, TRAF6, TLR6, TLR2, TLR7, TICAM1 | 4.34510E-15 |
| lupus nephritis | 438 | 10 | 2 | TLR3, TLR4, TLR9, TLR5, TRAF3, MYD88, TRAF6, IRAK4, TLR2, TLR7 | 8.96976E-15 |
| neutrophilic inflammation | 279 | 9 | 3 | TLR3, TLR4, TLR9, MYD88, TLR8, TLR6, TLR2, TLR7, TICAM1 | 1.42359E-14 |
| Brucella infection | 77 | 7 | 8 | TLR3, TLR4, TLR9, MYD88, TLR6, TLR2, TLR7 | 1.51854E-14 |
| low grade inflammation | 169 | 8 | 4 | TLR3, TLR4, TLR9, TLR5, MYD88, TRAF6, TLR8, TLR2 | 2.83711E-14 |

**Table 8**: shows the diseases targeted by entities in the gene set.

**Table 8.** Presence in organs of Gene set. TLR1-10, MYD88, IRAK4,TRAF6,TRAF3, TICAM1(TRIF)

| Protein entities related to | Total # of Neighbors | Overlap | Percent Overlap | Overlapping Entities | p-value |
| --- | --- | --- | --- | --- | --- |
| Testis | 10388 | 13 | 0 | TLR3, TLR4, TLR10, TLR5, MYD88, IRAK4, TLR9, TLR1, TLR8, TLR6, TLR2, TLR7, TICAM1 | 0.00000E+00 |
| cerebral cortex | 8885 | 12 | 0 | TLR3, TLR4, TLR5, MYD88, IRAK4, TLR9, TRAF3, TRAF6, TLR8, TLR6, TLR2, TLR7 | 0.00000E+00 |
| brain | 11791 | 15 | 0 | TLR3, TLR4, TLR10, TLR5, MYD88, IRAK4, TLR9, TRAF3, TRAF6, TLR1, TLR8, TLR6, TLR2, TLR7, TICAM1 | 0.00000E+00 |
| neuron | 9349 | 14 | 0 | TLR3, TLR4, TLR5, MYD88, IRAK4, TLR9, TRAF3, TRAF6, TLR1, TLR8, TLR6, TLR2, TLR7, TICAM1 | 0.00000E+00 |
| lung | 10058 | 15 | 0 | TLR3, TLR4, TLR10, TLR5, MYD88, IRAK4, TLR9, TRAF3, TRAF6, TLR1, TLR8, TLR6, TLR2, TLR7, TICAM1 | 0.00000E+00 |
| kidney | 10916 | 14 | 0 | TLR3, TLR4, TLR5, MYD88, IRAK4, TLR9, TRAF3, TRAF6, TLR1, TLR8, TLR6, TLR2, TLR7, TICAM1 | 0.00000E+00 |
| liver | 10017 | 15 | 0 | TLR3, TLR4, TLR10, TLR5, MYD88, IRAK4, TLR9, TRAF3, TRAF6, TLR1, TLR8, TLR6, TLR2, TLR7, TICAM1 | 0.00000E+00 |
| EPC | 84 | 11 | 12 | TLR3, TLR4, TLR9, TLR10, TLR5, MYD88, TLR1, TLR8, TLR6, TLR2, TLR7 | 5.27370E-20 |
| OKF6 | 46 | 9 | 19 | TLR3, TLR4, TLR9, TLR5, TLR1, TLR8, TLR6, TLR2, TLR7 | 7.73300E-18 |
| IPEC-J2 | 234 | 11 | 4 | TLR3, TLR4, TLR9, TLR10, TLR5, MYD88, TLR1, TLR8, TLR6, TLR2, TLR7 | 6.04630E-15 |

**Table 9**: shows expression of the genes in various organs and table 9shows the key pathological proce4sses involved.

**Table 9.**
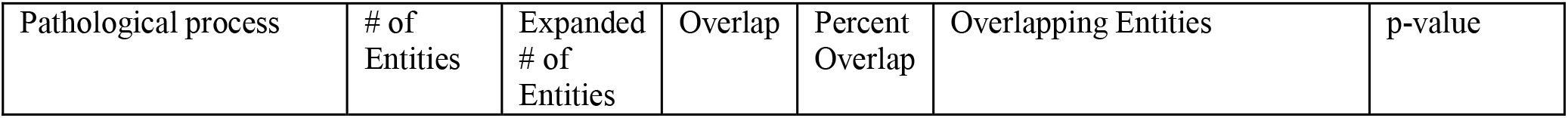
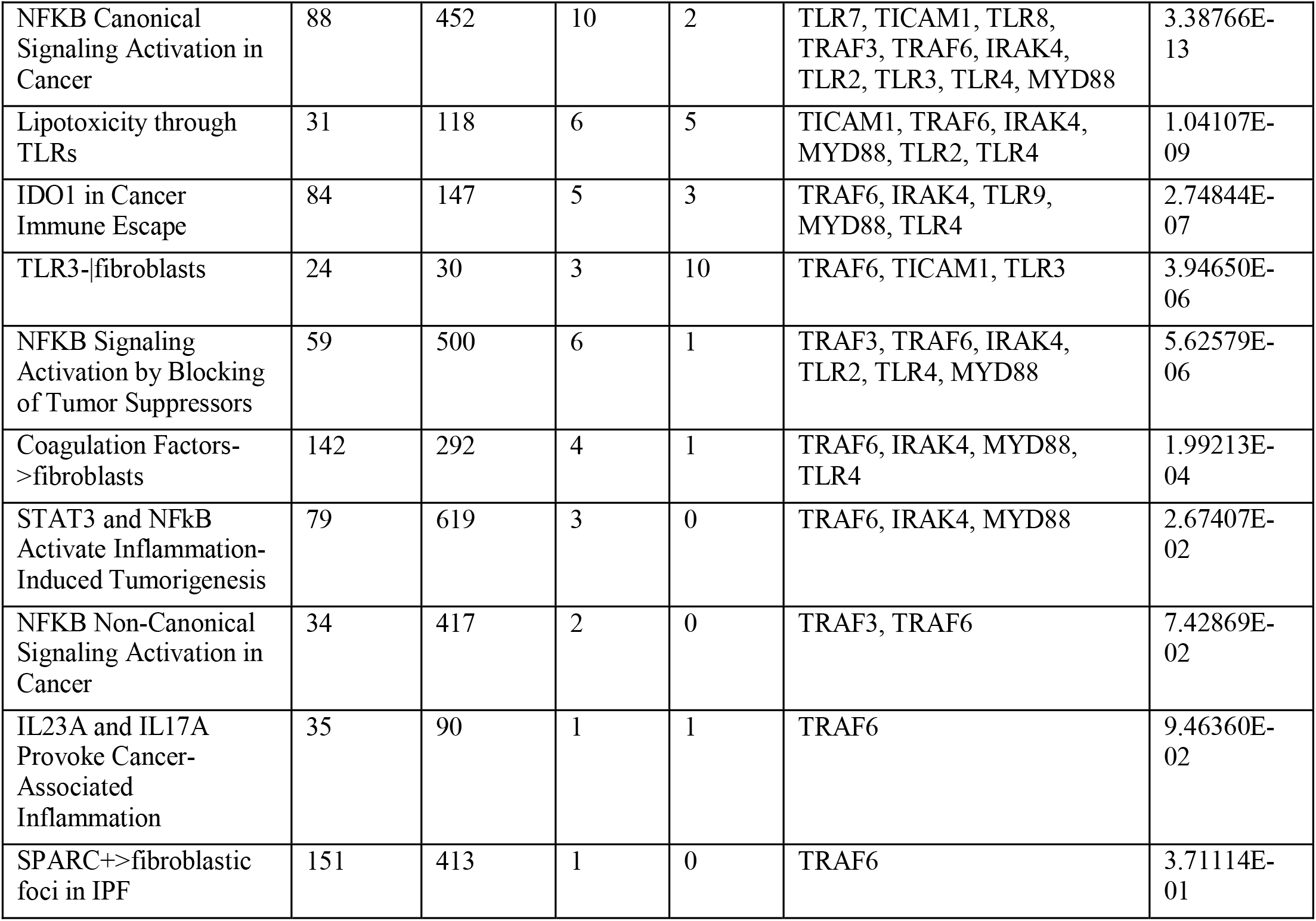
Pathological process.

TLR4 mRNA has been detected in the testis, indicating a potential role in immune responses within this organ. TLR4 expression has been detected in various brain regions, such as the hypothalamus, hippocampus, cortex, and cerebellum. TLR4 expression varies among species but occurs in different animal species (82).

TLR4 mRNA is found in the lungs of various species, such as pigs and rabbits. TLR2 and TLR4 are implicated in renal diseases such as acute kidney injury (AKI) and lupus nephritis (LN). TLR4 expression levels vary among mouse strains and contribute to producing pro- and anti-inflammatory cytokines (83).

TLRs such as TLR2 and TLR4 play a role in the development and progression of renal diseases by influencing inflammatory responses (83). Inflammatory responses: TLR activation is crucial for initiating inflammatory responses by recognizing microbial or endogenous ligands (13). Organ-specific effects: The differential expression of TLRs across organs can impact immune responses and disease outcomes. For example, variations in TLR expression on endothelial cells can influence inflammatory disorders (84). Their expression patterns in organs such as the testis, brain, lung, kidney, and liver suggest their involvement in immune responses and disease processes specific to each organ and provide insights into their diverse roles beyond traditional immune function.

### Summary

TLRs initiate an immune response by recognizing and responding to a vast array of PAMPs, including flagellin, peptidoglycans, bacterial lipopolysaccharides, lipoproteins, peptidoglycans, and nucleic acids derived from bacteria and viruses. This is shown (figure 4) by inputting genes set in KEGG Pathway (85,86). This is shown in figure 4 and it illustrates how Metadichol activates multiple pathways to inhibit various bacteria and viruses The TLR network regulates numerous biological processes, and its investigation is crucial for comprehending the immune response in various species. The ecological significance of TLRs across species has been established by recent genome surveys that have revealed their presence in nonmammalian organisms (87). This finding further supports the notion that TLRs have undergone evolutionary conservation. Metadichol induces an inverted U response in all proteins, including MYD88, and is specific enough to downregulate TLR4, which has been implicated in numerous maladies. Metadichol is the only known molecule capable of expressing the complete family of TLR receptors. Furthermore, they express neuronal transcription factors, Yamanaka factors, nuclear receptors, and cardiovascular transcription factors (88-92). These factors are also detected in fibroblasts at a minimal concentration of one picogram per milliliter. By modulating a family of crucial transcription factors, it is capable of alleviating a multitude of diseases (93). Thus far, the objective of drug research has been to develop exceptionally selective molecules that specifically target a single biomolecule that is implicated in the pathogenesis of a given disease (94). Such a strategy may prove beneficial for diseases whose mechanisms are well defined. Although thousands of diseases affect humans, a single-target, single-drug strategy has proven ineffective (95). There is a need for multitarget pharmaceuticals that exert their effects on numerous genes or gene families, as diseases are influenced by a multitude of factors (96-97). The actions of metadichol constitute a network and multitargeting approach (98). By selectively targeting telomerase, sirtuins, TLRs, and nuclear receptors, among other factors, metadichol enables the exploration of potential therapeutic interventions in the fields of anti-aging and cell reprogramming and the elucidation of the molecular mechanisms that govern cellular function and aging. A solution cannot be found with single-target agents for most diseases. Metadichol® introduces a novel paradigm for advancements in pharmaceutical research and development. Due to its nontoxic characteristics (99-101), it has the potential to significantly contribute to the resolution of many human disease conditions for which no viable solution has yet been found.

**Figure 4.**
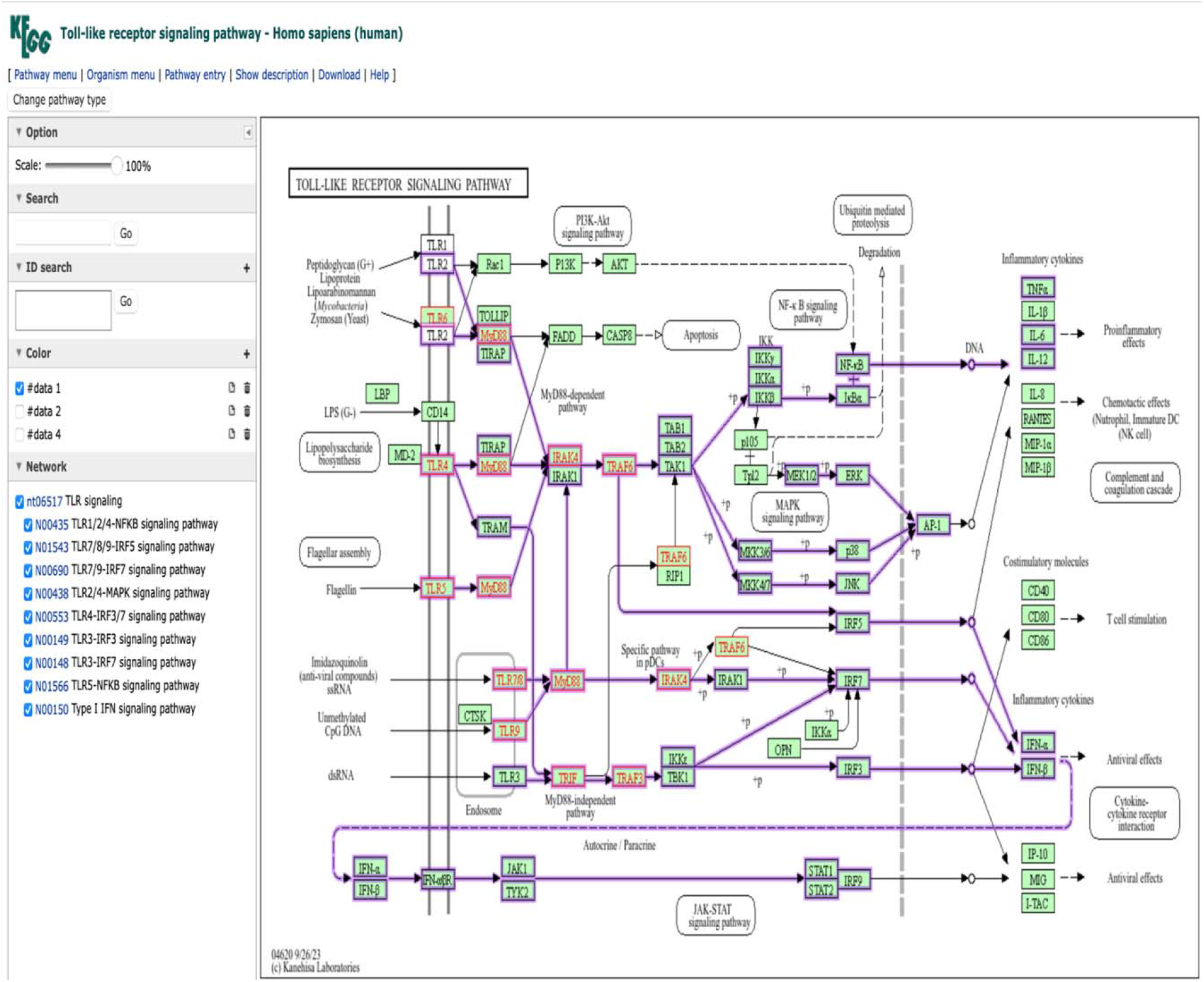
Metadichol expressed genes and Toll receptor Signaling pathway

## Supporting information

Raw Experimental Data

## Data availability

**A**ll the raw data are presented in the manuscript and in the supplementary materials.

## Supplementary material

Raw Data: Q-RT□PCR

