## Supplementary material for "Metadichol® induced expression of the TLR family of receptors in PBMCs": Raw Experimental Data

### Raw Data og Fold changes by Q-Rt-pcr

#### Evaluation of effect of Metadichol on gene expression of Toll like receptor using Human peripheral mononuclear blood cells (PBMCs)

### Evaluation of effect of Metadichol on gene expression of Toll like receptor using Human peripheral mononuclear blood cells (PBMCs)

#### 1] Cellular markers

1] TLR 1

2] TLR 2

3] TLR 3

4] TLR 4

5] TLR 5

6] TLR 6

7] TLR 7

8] TLR 8

9] TLR 9

10] TLR 10

11] MYD88

12] IRAK4

13] TRAF3

14] TRAF6

15] TRIF

#### 2] Chemicals/ reagents used

1] Sterile workplace, tips, pipettes, centrifuge tubes, 6-well plate etc.

2] Fresh Human Blood

3] Histopaque-1077(Sigma –Aldrich, Cat No.10771)

4] RPMI or DMEM Medium

5] Fetal bovine serum (FBS)

6] 0.05% trypsin

7] RNase free environment or workstation

8] Autoclaved deionised water

9] TRIzol reagent (Life technologies, Cat. No. 15596026)

10] Chloroform

11] Chilled isopropanol

12] Chilled 70% ethanol

13] Ice cold Phopshate buffered saline (PBS, pH 7.4)

14] Autoclaved, oven dried 1.5 and 2 ml micro-centrifuge tubes and PCR tubes

15] cDNA synthesis kit or Prime script RT Reagent kit (TAKARA, Cat. No. RR037A)

16] SyBR Green I (PCR Biosystems, Cat. No. PB20.15)

17] Dimethyl Sulphoxide (DMSO)

18] Target specific forward and reverse primer etc.

#### 3] Cell line and cell condition

#### 1. Isolation of Human WBCs

**Preparation of the blood sample-** Fresh human blood was collected in EDTA containing tubes, and then fresh blood was diluted with PBS in 1:1 proportion and mixed by inverting the tube.

**Isolation of Mononuclear Cells-** In the 15 ml centrifuge tube, 5 ml of Histopaque-1077 was added to that 5 ml of Prepared blood was layered on histopaque slowly from edge of the tube without disturbing the histopaque layer. Then tubes were centrifuged at 400 X g for exactly 30 mins at room temperature with brake off settings. After centrifugation, upper layer was discarded with Pasteur pipette without disturbing interphase layer. The interphase layer was carefully transferred to clean centrifuge tube. Cells will be washed with 1X PBS and again centrifuged at 250 X g for 10 mins. (2X). After centrifugation, supernatant was discarded and pellet will be collected in RPMI media supplemented with 10% FBS. Cells were counted and viability was checked with Hemo-cytometer.

**Cell maintenance and seeding-** Cell density at 1 X $10^6$ cells/ml of media were prepared and seeded into 6 well plates and incubated for 24 hrs at 37°C with 5% $CO_2$. Post 24 hrs of seeding, the media was carefully removed and the cells were treated with respective test samples (Concentration will be selected on basis of MTT experiment) and incubated for 24 hrs at 37°C in $CO_2$ incubator.

**Table 1: Treatment concentrations**

| Sr. No | **Cell line** | **Sample name** | **Treatment details** |
|---|---|---|---|
| 1 | Human PBMC | Metadichol | Control |
| | | | 1 pg |
| | | | 100 pg |
| | | | 1 ng |
| | | | 100ng |

**Sample Preparation and RNA Isolation**

Treated cells were dissociated and rinsed with sterile 1X PBS and centrifuged. The supernatant was decanted and 0.1 ml of TRIzol was added and gently mixed by inversion for 1 min. Samples were allowed to stand for 10 minutes at room temperature. To this 0.75 ml chloroform was added per 0.1 ml of TRIzol used. The contents were vortexed for 15 seconds. The tube was allowed to stand at room temperature for 5 mins. The resulting mixture was centrifuged at 12,000 rpm for 15 mins at 4˚C. Upper aqueous phase was collected to a new sterile micro-centrifuge tube to which 0.25 ml of isopropanol was added and gently mixed by inverting the contents for 30 seconds and incubated at -20˚C for 20 minutes. The contents were centrifuged at 12,000 rpm for 10 minutes at 4˚C. Supernatant was discarded and the RNA pellet was washed by adding 0.25 ml of 70% ethanol. The RNA mixture was centrifuged at 12,000 rpm at 4˚C. Supernatant was carefully discarded and the pellet was air dried. The RNA pellet was then re-suspended in 20 µl of DEPC treated water. Total RNA yield was quantified using Spectra drop (Spectramax i3x, Molecular devices, USA).

**Table 2: Total RNA yield**

| | **Test concentrations** | | | | |
|---|---|---|---|---|---|
| **RNA yield (ng/µl)** | **0** | **1 pg/ ml** | **100 pg/ ml** | **1 ng/ ml** | **100 ng/ ml** |
| **Human PBMC's** | 438.00 | 304.64 | 161.92 | 376.24 | 382.88 |

#### qPCR analysis

##### cDNA synthesis

The cDNA was synthesized from 500 ng of RNA using the cDNA synthesis kit from Prime script RT reagent kit (TAKARA) with oligo dT primer according to the manufacturer's instructions. The reaction volume was set to 20 µl and cDNA synthesis was performed at 50 ºC for 30 min, followed by RT inactivation at 85ºC for 5 min using applied biosystems, Veritii. The cDNA was further used for real time PCR analysis.

##### Primers and qPCR analysis

The PCR mixture (final volume of 20 µl) contained 1.4 µl of cDNA, 10 µL of SyBr green Master mix and 1 µM of respective complementary forward and reverse primers specific for respective target genes. The reaction was carried out with enzyme activation at 95ºC for 2 minutes followed by 2 step reaction with initial denaturation and annealing cum extension step at 95°C for 5 seconds, annealing for 30 seconds at appropriate respective temperature amplified for 39 cycles followed by secondary denaturation at 95 °C for 5 seconds, 1 cycle with melt curve capture step ranging from

65°C to 95°C for 5 secs each. The obtained results was analysed and fold expression or regulation was calculated

**Table 3: Primer details**

| Sr. no. | Primer | Sequence | Amplicon size | Annealing temperature |
|---|---|---|---|---|
| 1 | GAPDH | GTCTCCTCTGACTTCAACAGCG | 186 | 60 |
| | | ACCACCCTGTTGCTGTAGCCAA | | |
| 2 | TLR 1 | CAGCGATGTGTTCGGTTTTCCG | 145 | 67 |
| | | GATGGGCAAAGCATGTGGACCA | | |
| 3 | TLR 2 | CTTCACTCAGGAGCAGCAAGCA | 110 | 67 |
| | | ACACCAGTGCTGTCCTGTGACA | | |
| 4 | TLR 3 | CACCATTCCAGCCTCTTCGT | 157 | 65 |
| | | CAGGGTTTGCGTGTTTCCAG | | |
| 5 | TLR 4 | CCCTGAGGCATTTAGGCAGCTA | 125 | 65 |
| | | AGGTAGAGAGGTGGCTTAGGCT | | |
| 6 | TLR 5 | CCTTACAGCGAACCTCATCCAC | 129 | 65 |
| | | TCCACTACAGGAGGAGAAGCGA | | |
| 7 | TLR 6 | ACTGACCTTCCTGGATGTGGCA | 113 | 67 |
| | | TGACCTCATCTTCTGGCAGCTC | | |
| 8 | TLR 7 | CTTTGGACCTCAGCCACAACCA | 141 | 67 |
| | | CGCAACTGGAAGGCATCTTGTAG | | |
| 9 | TLR 8 | ACTCCAGCAGTTTCCTCGTCTC | 144 | 65 |
| | | AAAGCCAGAGGGTAGGTGGGAA | | |
| 10 | TLR 9 | CTGCCTTCCTACCCTGTGAG | 138 | 67 |
| | | GGATGCGGTTGGAGGACAA | | |
| 11 | TLR 10 | GGTTCTTTTGCGTGATGGAATC | 164 | 65 |
| | | GGTCGTCCCAGAGTAAATCAAC | | |
| 12 | MYD88 | CACCACACTTGATGACCCCC | 215 | 62 |
| | | TCCGGCGGCACCTCTTTT | | |
| 13 | IRAK4 | CATTCCCCGCCTTAATGCCT | 157 | 67 |
| | | AGCTCTGTCAAAGGAACCCA | | |
| 14 | TRAF3 | GCGTGTCAAGAGAGCATCGTT | 133 | 59 |
| | | GCAGATGTCCCAGCATTAACT | | |
| 15 | TRAF6 | CAATGCCAGCGTCCCTTCCAAA | 141 | 65 |
| | | CCAAAGGACAGTTCTGGTCATGG | | |
| 16 | TRIF | CCTGGAATCATCATCGGAACAG | 193 | 65 |
| | | TGAGTGGTCTATGGCGTCCT | | |

### Results-

#### 1. GAPDH

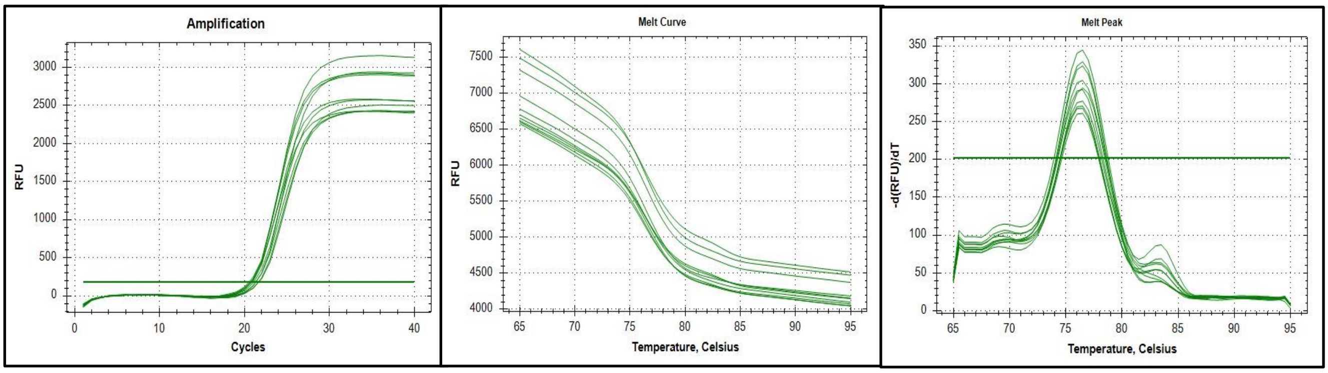


**Fig 1: Amplification curve, Melt curve and Melt peak of GAPDH gene**

#### 2. TLR 1

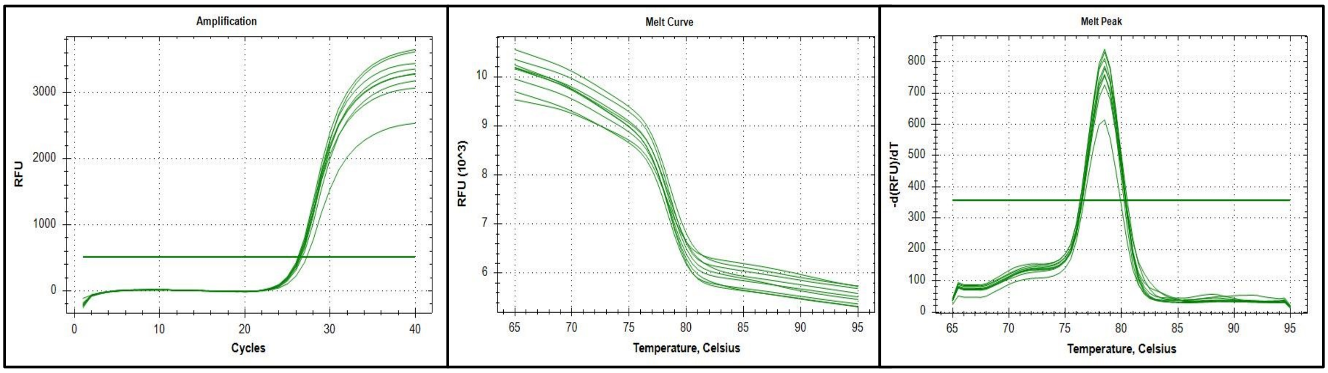


**Fig 2: Amplification curve, Melt curve and Melt peak of TLR 1 gene**

#### 3. TLR 2

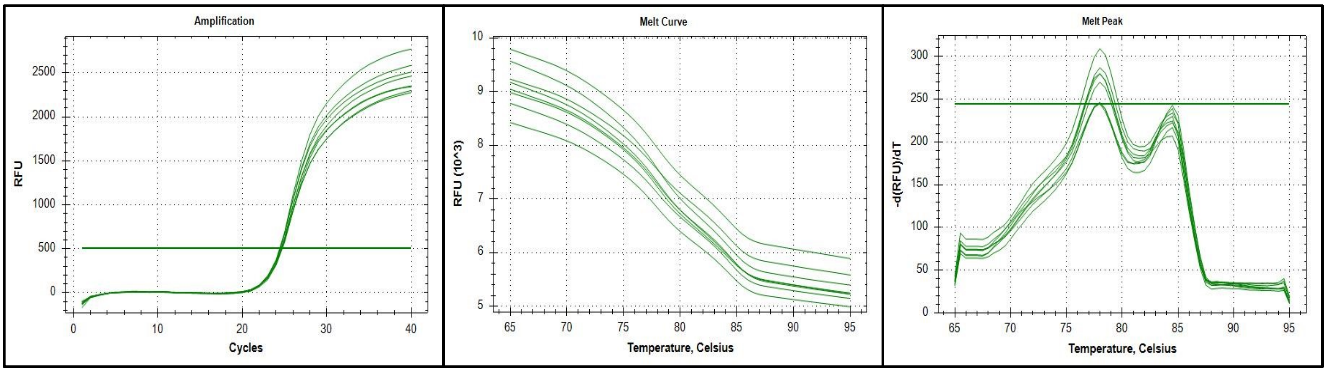


**Fig 3: Amplification curve, Melt curve and Melt peak of TLR 2 gene**

#### 4. TLR 3

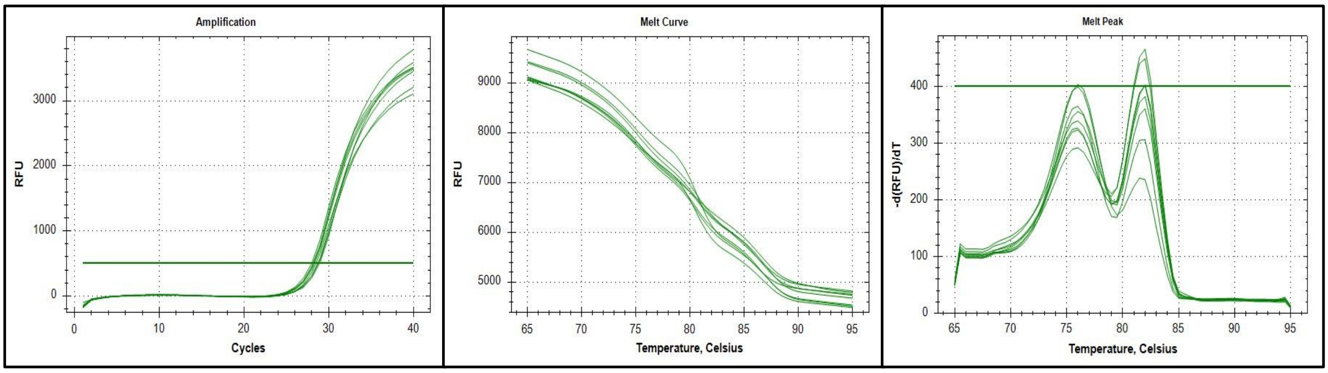


**4: Amplification curve, Melt curve and Melt peak of TLR 3 gene**

#### 5. TLR 4

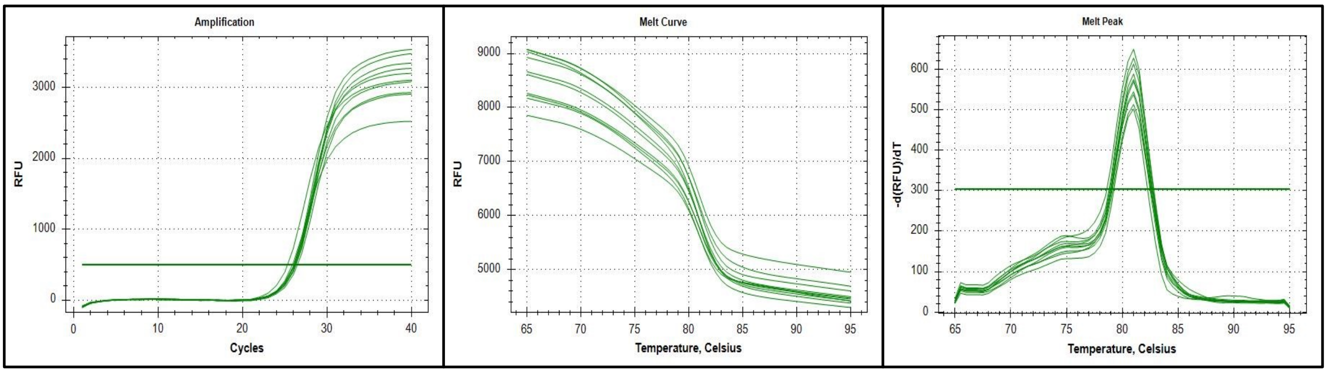


**Fig 5: Amplification curve, Melt curve and Melt peak of TLR 4 gene**

#### 6. TLR 5

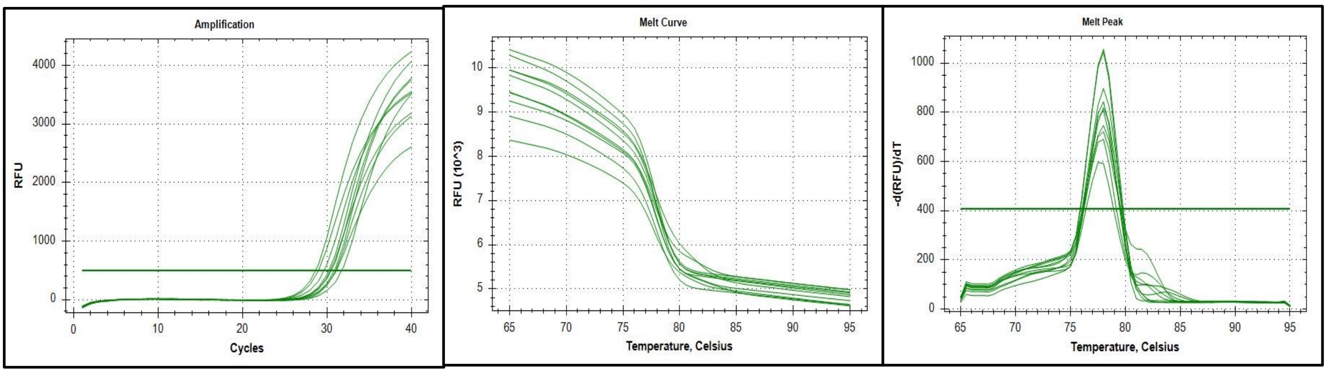


**Fig 6: Amplification curve, Melt curve and Melt peak of TLR 5 gene**

7. **TLR 6**

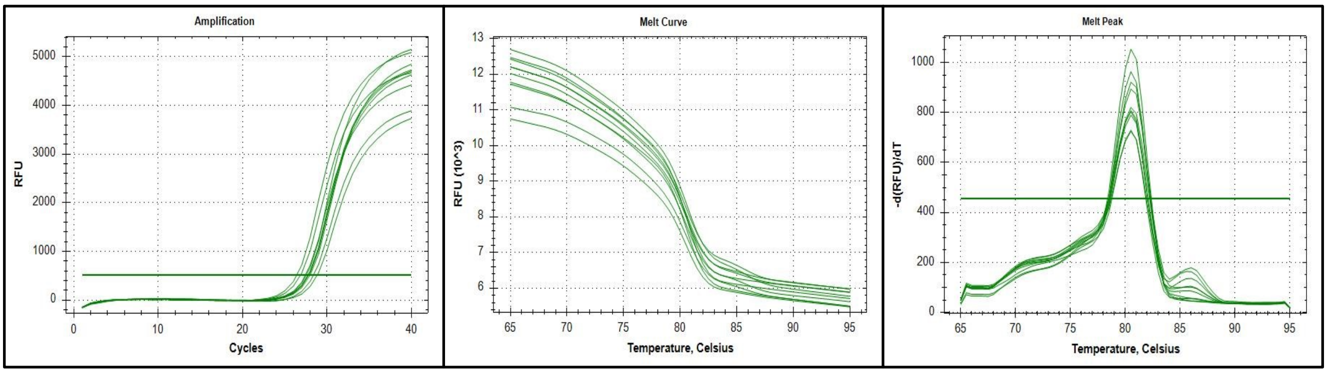


**Fig 7: Amplification curve, Melt curve and Melt peak of TLR 6 gene**

8. **TLR 7**

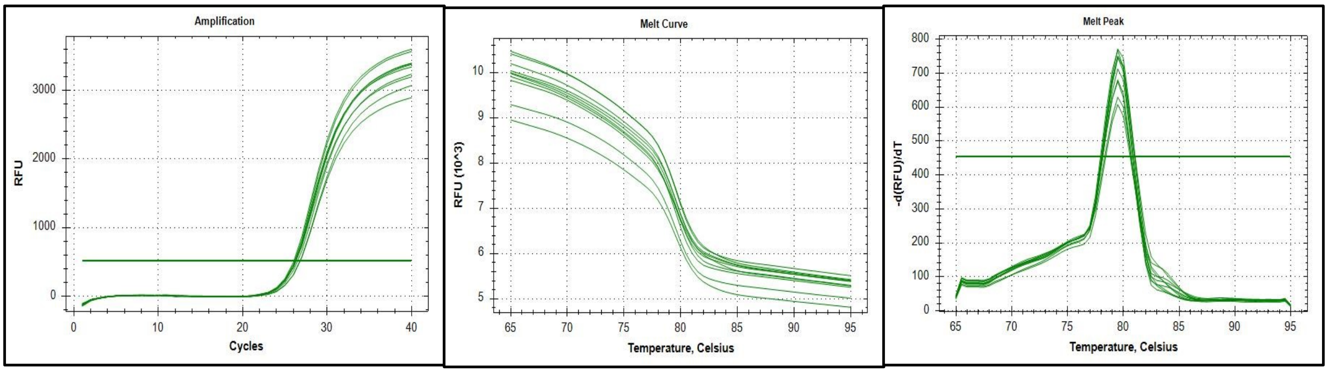


**Fig 8: Amplification curve, Melt curve and Melt peak of TLR 7 gene**

9. **TLR 8**

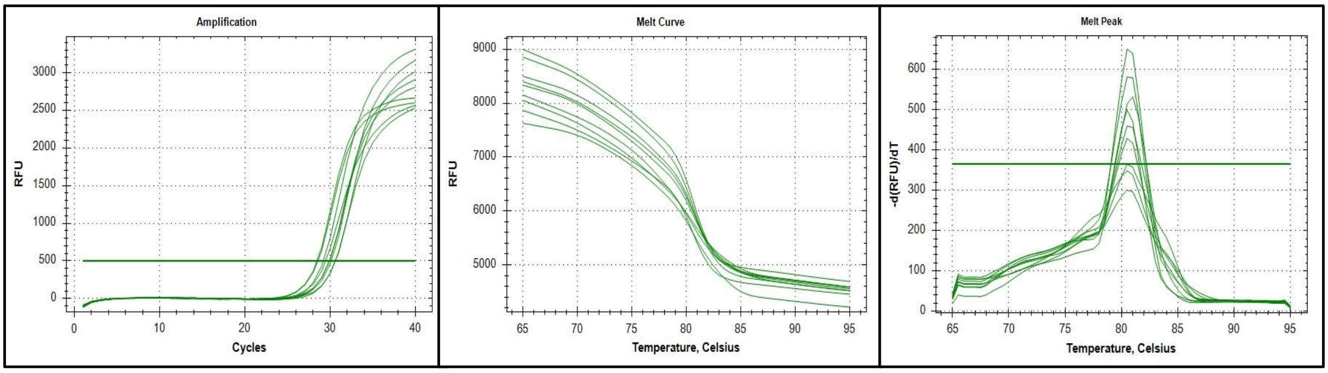


**Fig 9: Amplification curve, Melt curve and Melt peak of TLR 8 gene**

#### 10. TLR 9

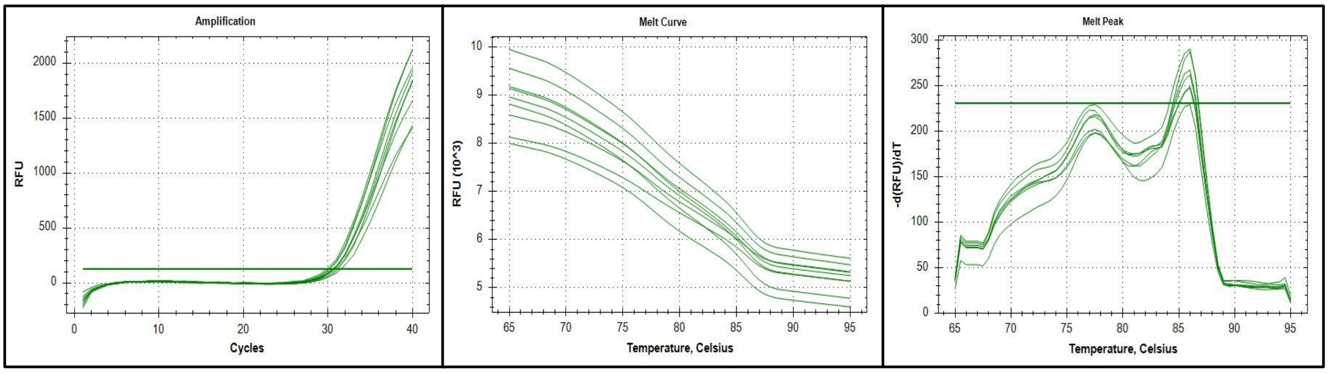


**Fig 10: Amplification curve, Melt curve and Melt peak of TLR 9 gene**

#### 11. TLR 10

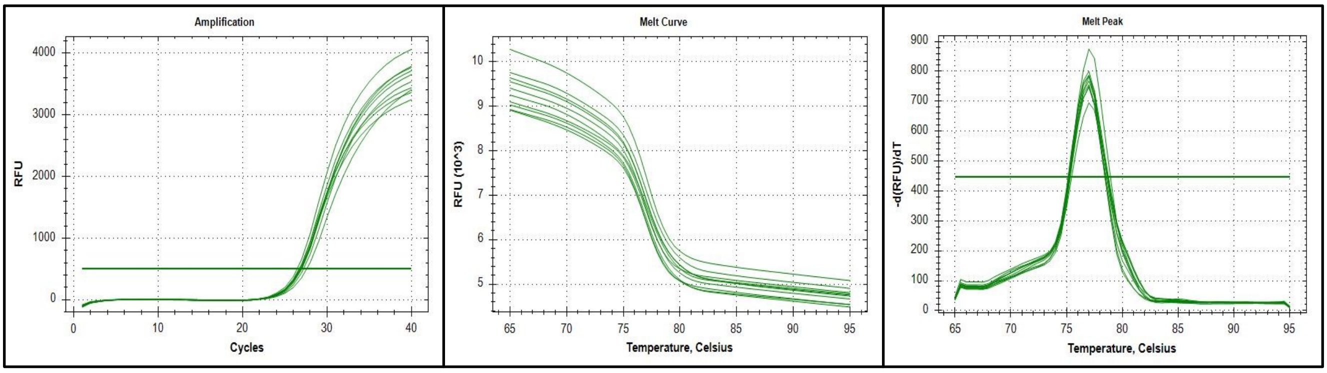


**Fig 11: Amplification curve, Melt curve and Melt peak of TLR 10 gene**

#### 12. MYD88

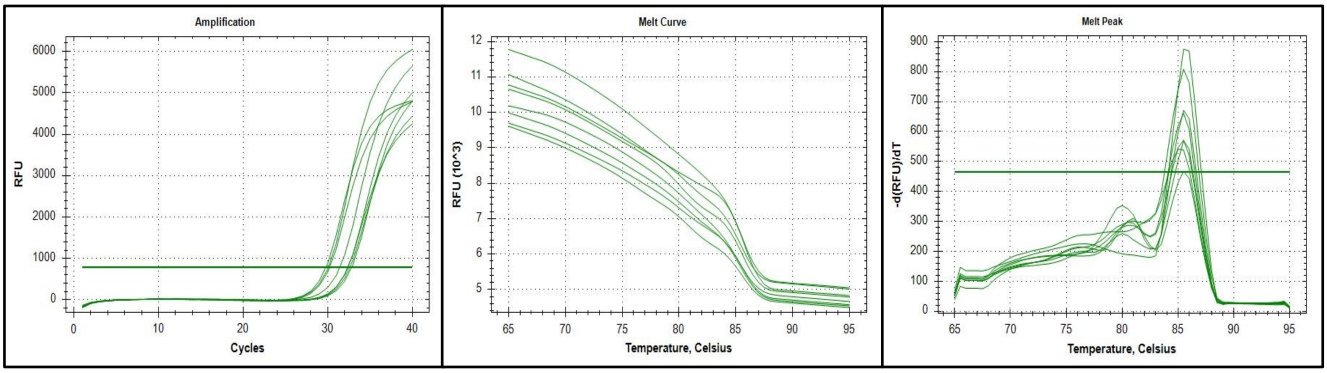


**Fig 12: Amplification curve, Melt curve and Melt peak of MYD88 gene**

#### 13. IRAK4

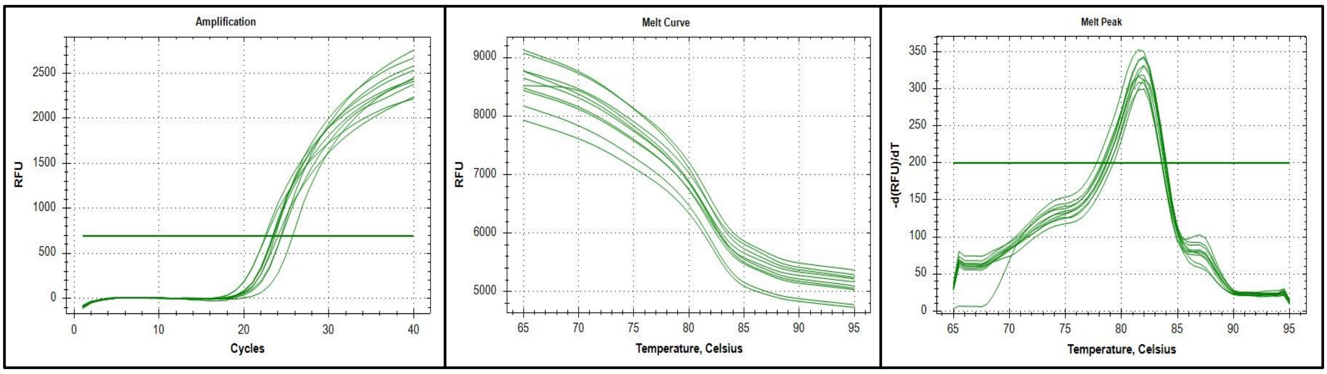


**Fig 13: Amplification curve, Melt curve and Melt peak IRAK4 gene**

#### 14. TRAF3

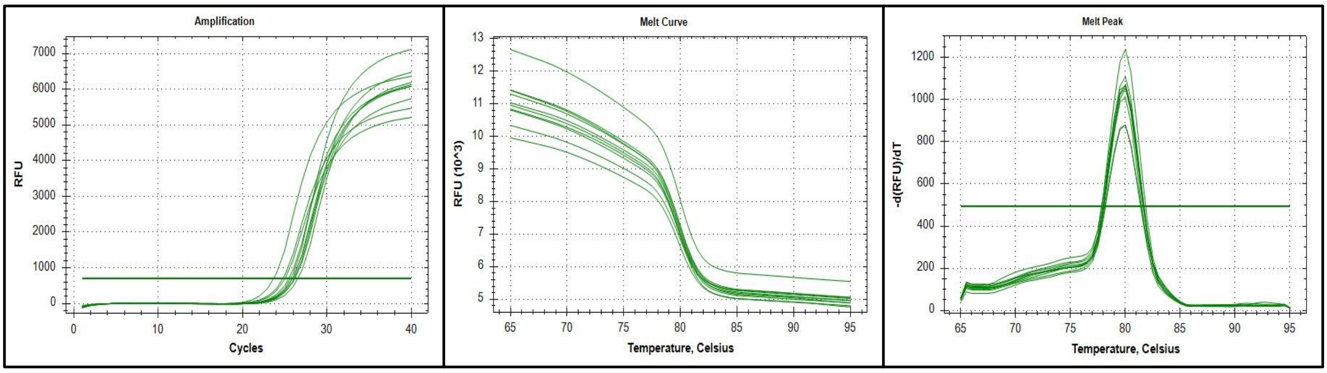


**Fig 14: Amplification curve, Melt curve and Melt peak TRAF3 gene**

#### 15. TRAF6

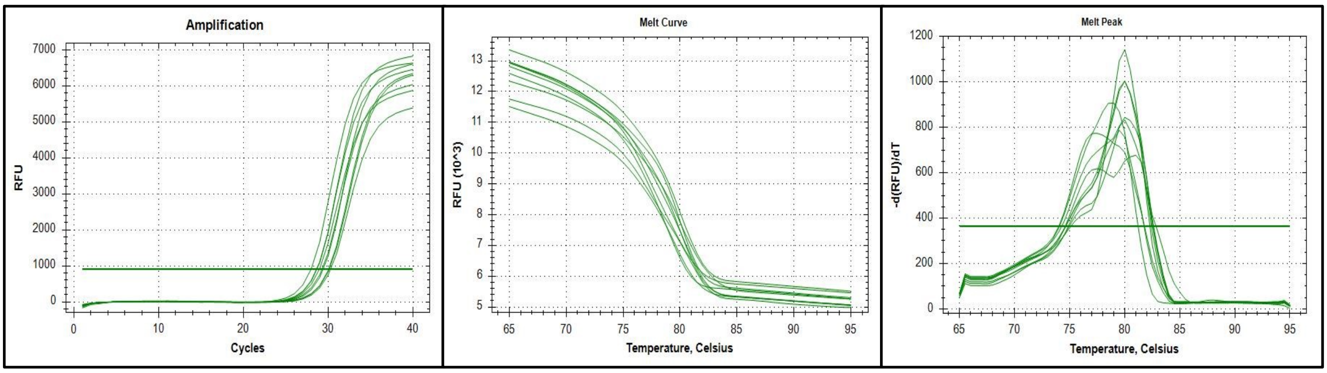


**Fig 15: Amplification curve, Melt curve and Melt peak TRAF6 gene**

#### 16. TRIF

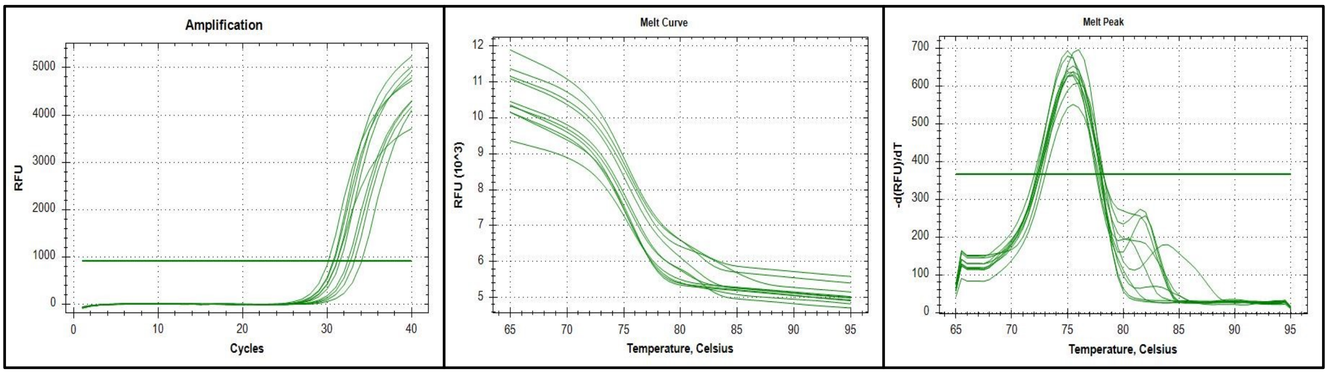


**Fig 16: Amplification curve, Melt curve and Melt peak TRIF gene**

**Table 4: Relative normalized expression of TLR 1 gene**

| Treatment Group | TLR 1 | GAPDH | ΔCq | ΔΔCq | Fold change 2^-ΔΔCq |
|---|---|---|---|---|---|
| Control | 27.12 | 20.66 | 6.46 | 0.00 | 1.00 |
| 1 pg | 26.33 | 20.86 | 5.47 | -0.99 | 1.99 |
| 100 pg | 26.44 | 21.47 | 4.97 | -1.49 | 2.81 |
| 1 ng | 26.34 | 21.09 | 5.25 | -1.21 | 2.31 |
| 100 ng | 26.56 | 21.03 | 5.53 | -0.93 | 1.91 |

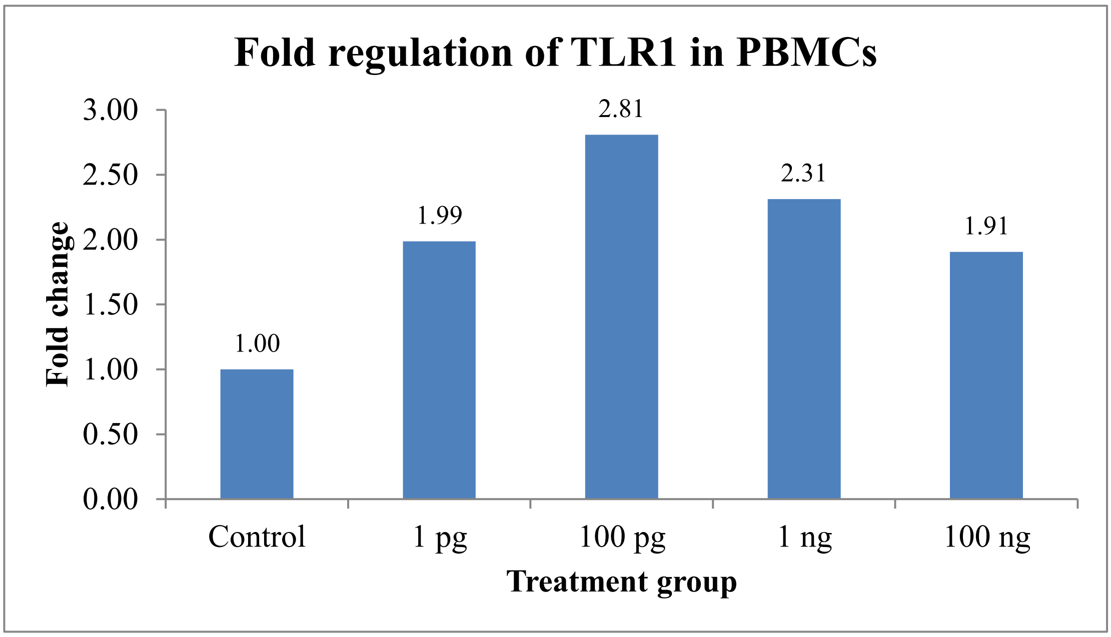


**Fig 17: Relative normalized expression of TLR 1 gene**

**Table 5: Relative normalized expression of TLR 2 gene**

| Treatment Group | TLR 2 | GAPDH | ΔCq | ΔΔCq | Fold change 2^-ΔΔCq |
|---|---|---|---|---|---|
| Control | 25.06 | 20.66 | 4.40 | 0.00 | 1.00 |
| 1 pg | 24.48 | 20.86 | 3.62 | -0.78 | 1.72 |
| 100 pg | 24.62 | 21.47 | 3.15 | -1.25 | 2.38 |
| 1 ng | 24.45 | 21.09 | 3.36 | -1.04 | 2.06 |
| 100 ng | 24.68 | 21.03 | 3.65 | -0.75 | 1.68 |

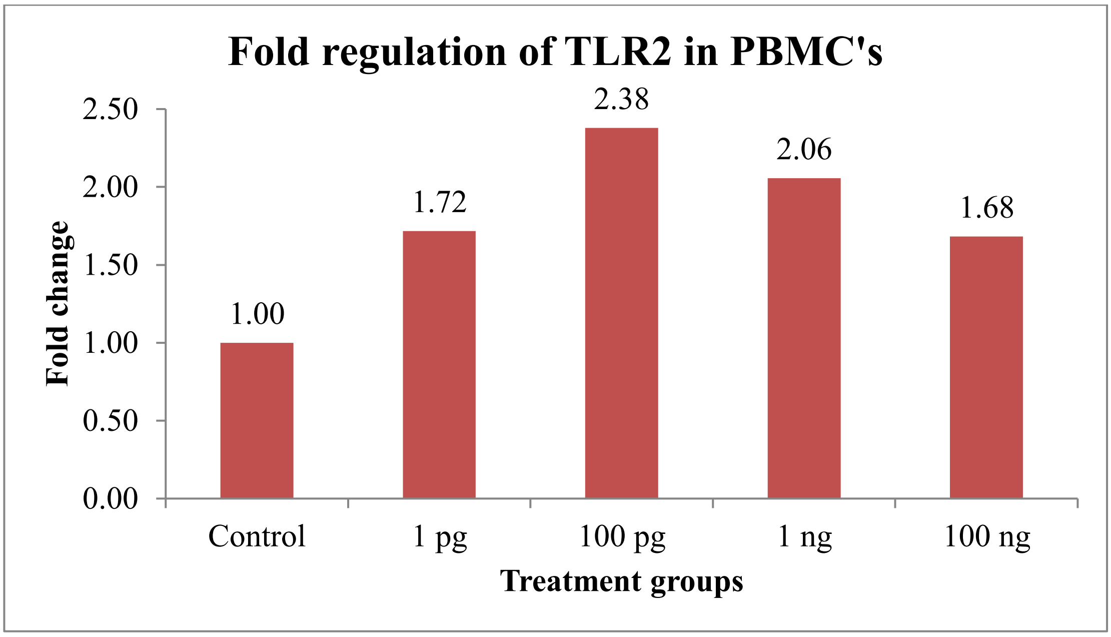


**Fig 18: Relative normalized expression of TLR 2 gene**

**Table 6: Relative normalized expression of TLR 3 gene**

| Treatment Group | TLR 3 | GAPDH | ΔCq | ΔΔCq | Fold change 2^-ΔΔCq |
|---|---|---|---|---|---|
| Control | 28.46 | 20.66 | 7.80 | 0.00 | 1.00 |
| 1 pg | 28.05 | 20.86 | 7.19 | -0.61 | 1.53 |
| 100 pg | 28.66 | 21.47 | 7.19 | -0.61 | 1.53 |
| 1 ng | 28.24 | 21.09 | 7.15 | -0.65 | 1.57 |
| 100 ng | 28.82 | 21.03 | 7.79 | -0.01 | 1.01 |

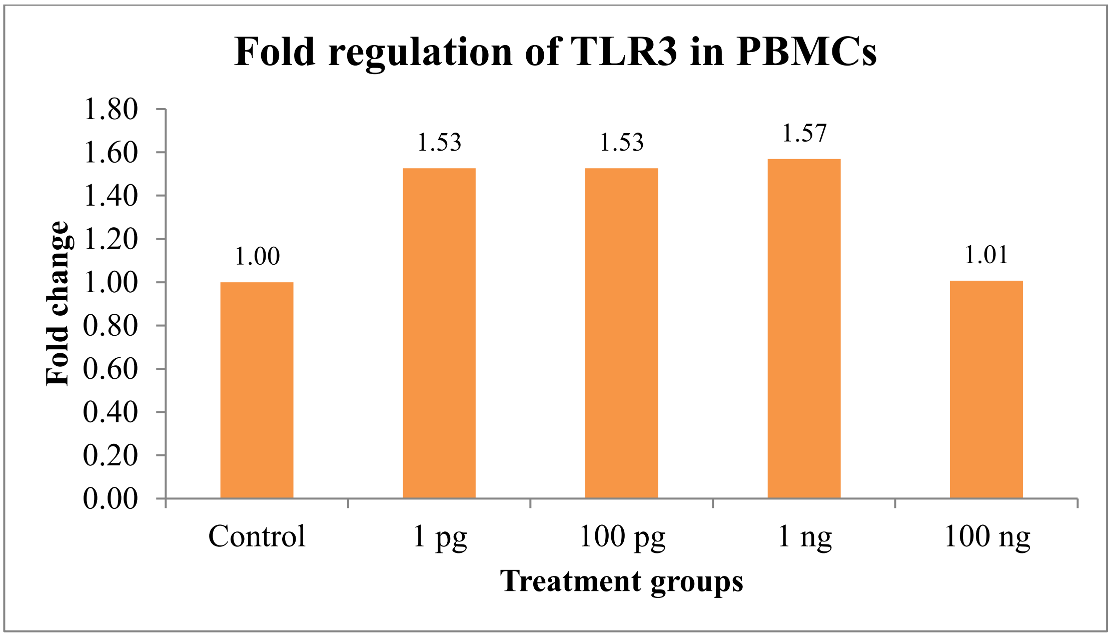


**Fig 19: Relative normalized expression of TLR 3 gene**

**Table 7: Relative normalized expression of TLR 4 gene**

| Treatment Group | TLR 4 | GAPDH | ΔCq | ΔΔCq | Fold change 2^-ΔΔCq |
|---|---|---|---|---|---|
| Control | 25.61 | 20.66 | 4.95 | 0 | 1.00 |
| 1 pg | 26.39 | 20.86 | 5.53 | 0.58 | 0.67 |
| 100 pg | 26.47 | 21.47 | 5.00 | 0.05 | 0.97 |
| 1 ng | 26.11 | 21.09 | 5.02 | 0.07 | 0.95 |
| 100 ng | 26.22 | 21.03 | 5.19 | 0.24 | 0.85 |

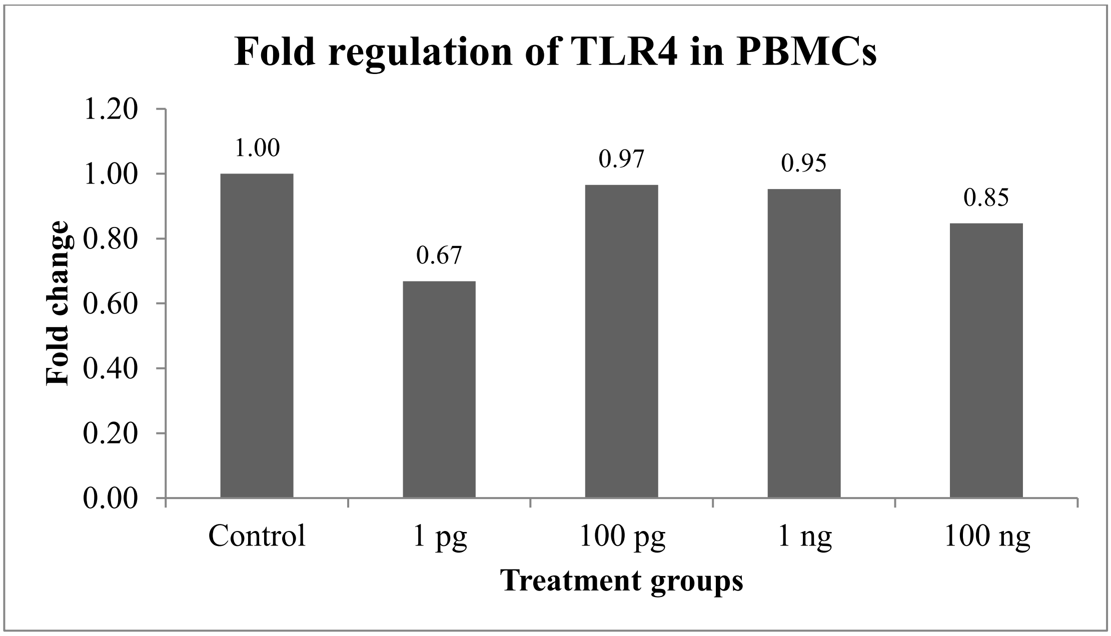


**Fig 20: Relative normalized expression of TLR 4 gene**

**Table 8: Relative normalized expression of TLR 5 gene**

| Treatment Group | TLR 5 | GAPDH | ΔCq | ΔΔCq | Fold change 2^-ΔΔCq |
|---|---|---|---|---|---|
| Control | 31.07 | 20.66 | 10.41 | 0.00 | 1.00 |
| 1 pg | 28.80 | 20.86 | 7.94 | -2.47 | 5.54 |
| 100 pg | 30.34 | 21.47 | 8.87 | -1.54 | 2.91 |
| 1 ng | 30.00 | 21.09 | 8.91 | -1.50 | 2.83 |
| 100 ng | 31.30 | 21.03 | 10.27 | -0.14 | 1.10 |

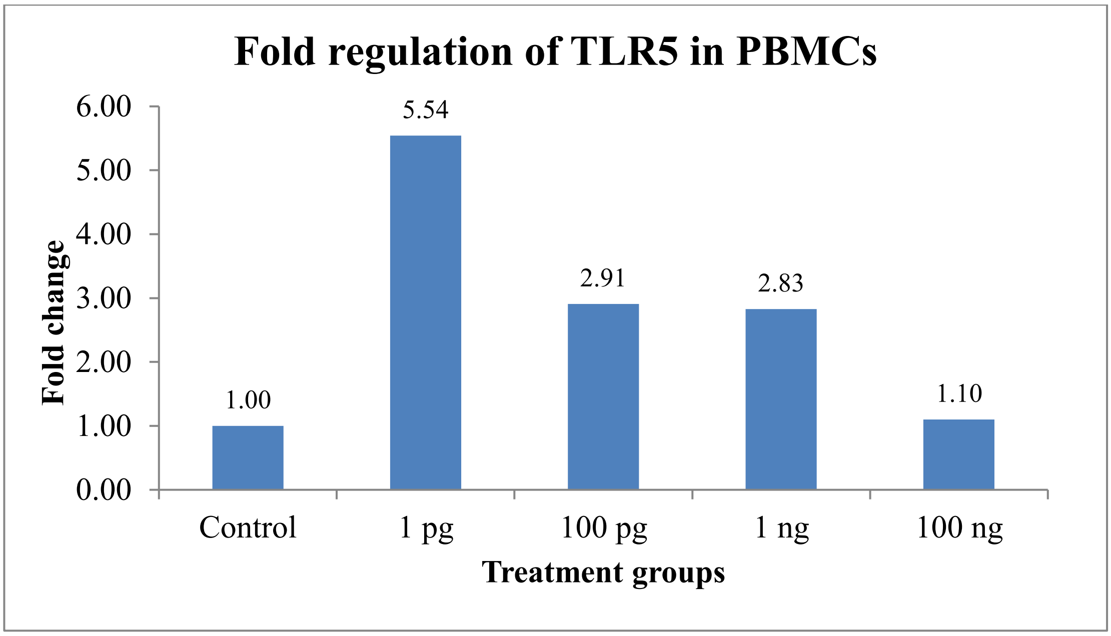


**Fig 21: Relative normalized expression of TLR 5 gene**

**Table 8: Relative normalized expression of TLR 6 gene**

| Treatment Group | TLR 6 | GAPDH | ΔCq | ΔΔCq | Fold change 2^-ΔΔCq |
|---|---|---|---|---|---|
| Control | 28.59 | 20.66 | 7.93 | 0.00 | 1.00 |
| 1 pg | 26.62 | 20.86 | 5.76 | -2.17 | 4.50 |
| 100 pg | 27.62 | 21.47 | 6.15 | -1.78 | 3.43 |
| 1 ng | 27.61 | 21.09 | 6.52 | -1.41 | 2.66 |
| 100 ng | 27.86 | 21.03 | 6.83 | -1.10 | 2.14 |

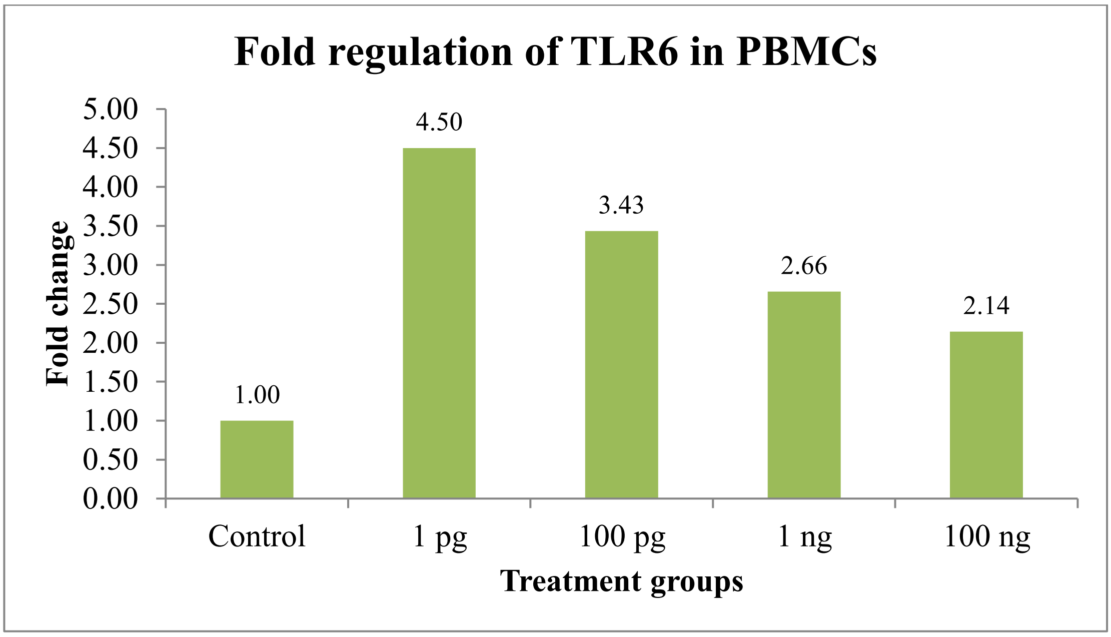


**Fig 22: Relative normalized expression of TLR 6 gene**

**Table 9: Relative normalized expression of TLR 7 gene**

| Treatment Group | TLR 7 | GAPDH | ΔCq | ΔΔCq | Fold change 2^-ΔΔCq |
|---|---|---|---|---|---|
| Control | 26.77 | 20.66 | 6.11 | 0.00 | 1.00 |
| 1 pg | 26.26 | 20.86 | 5.40 | -0.71 | 1.64 |
| 100 pg | 26.15 | 21.47 | 4.68 | -1.43 | 2.69 |
| 1 ng | 26.15 | 21.09 | 5.06 | -1.05 | 2.07 |
| 100 ng | 26.38 | 21.03 | 5.35 | -0.76 | 1.69 |

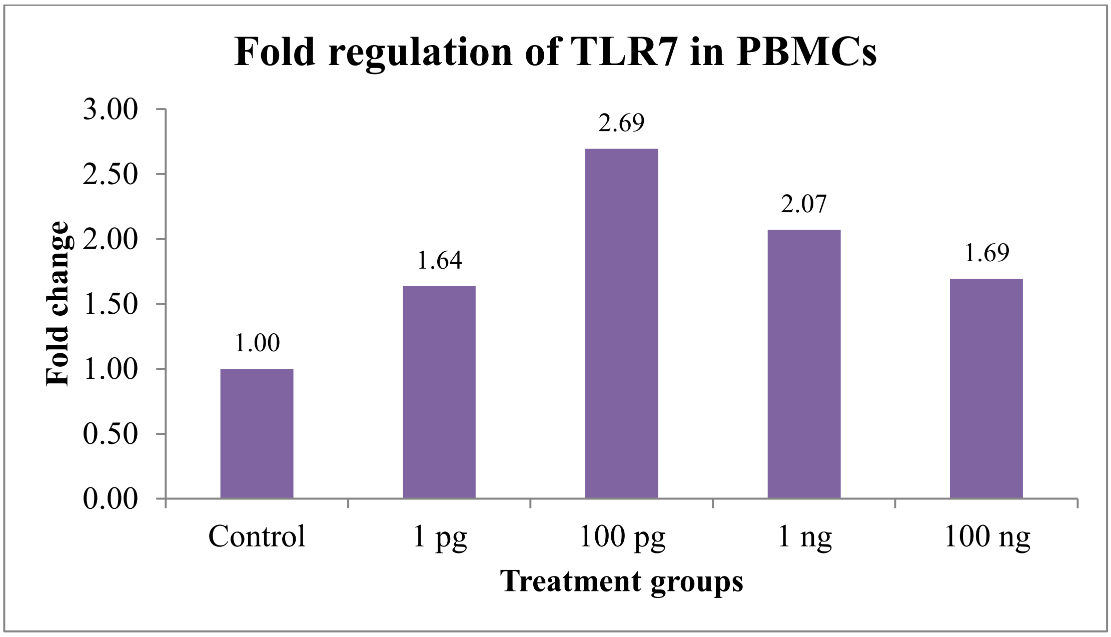


**Fig 23: Relative normalized expression of TLR 7 gene**

**Table 10: Relative normalized expression of TLR 8 gene**

| Treatment Group | TLR8 | GAPDH | ΔCq | ΔΔCq | Fold change 2^-ΔΔCq |
|---|---|---|---|---|---|
| Control | 30.40 | 20.66 | 9.74 | 0.00 | 1.00 |
| 1 pg | 29.32 | 20.86 | 8.46 | -1.28 | 2.43 |
| 100 pg | 28.50 | 21.47 | 7.03 | -2.71 | 6.54 |
| 1 ng | 29.92 | 21.09 | 8.83 | -0.91 | 1.88 |
| 100 ng | 30.36 | 21.03 | 9.33 | -0.41 | 1.33 |

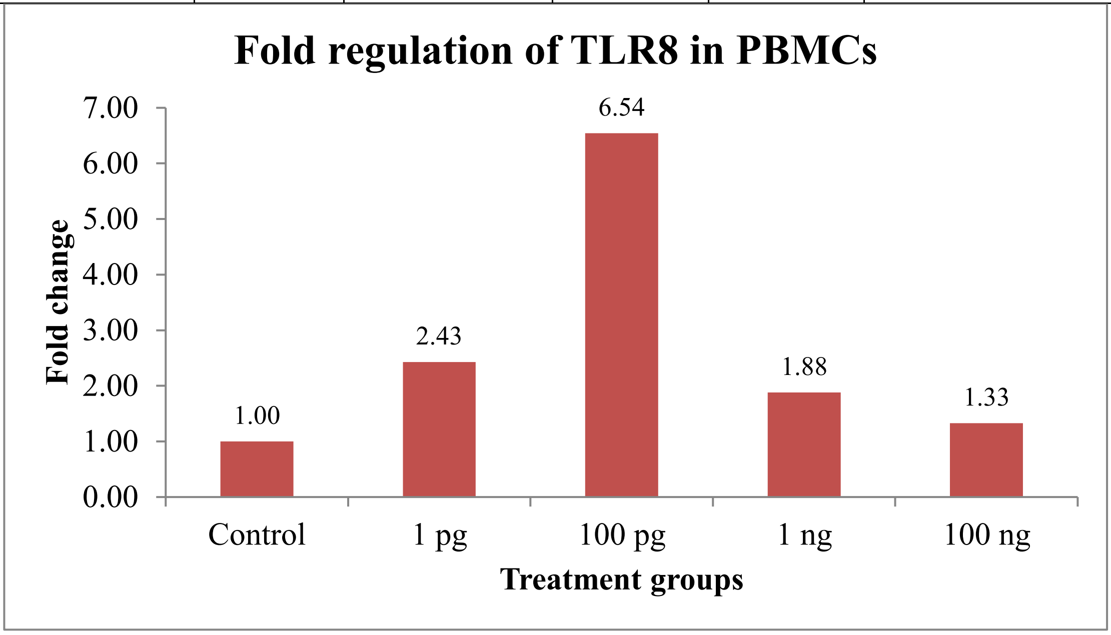


**Fig 24: Relative normalized expression of TLR 8 gene**

**Table 11: Relative normalized expression of TLR 9 gene**

| Treatment Group | TLR9 | GAPDH | ΔCq | ΔΔCq | Fold change 2^-ΔΔCq |
|---|---|---|---|---|---|
| Control | 30.43 | 20.66 | 9.77 | 0.00 | 1.00 |
| 1 pg | 30.45 | 20.86 | 9.59 | -0.18 | 1.13 |
| 100 pg | 30.34 | 21.47 | 8.87 | -0.90 | 1.87 |
| 1 ng | 30.32 | 21.09 | 9.23 | -0.54 | 1.45 |
| 100 ng | 30.66 | 21.03 | 9.63 | -0.14 | 1.10 |

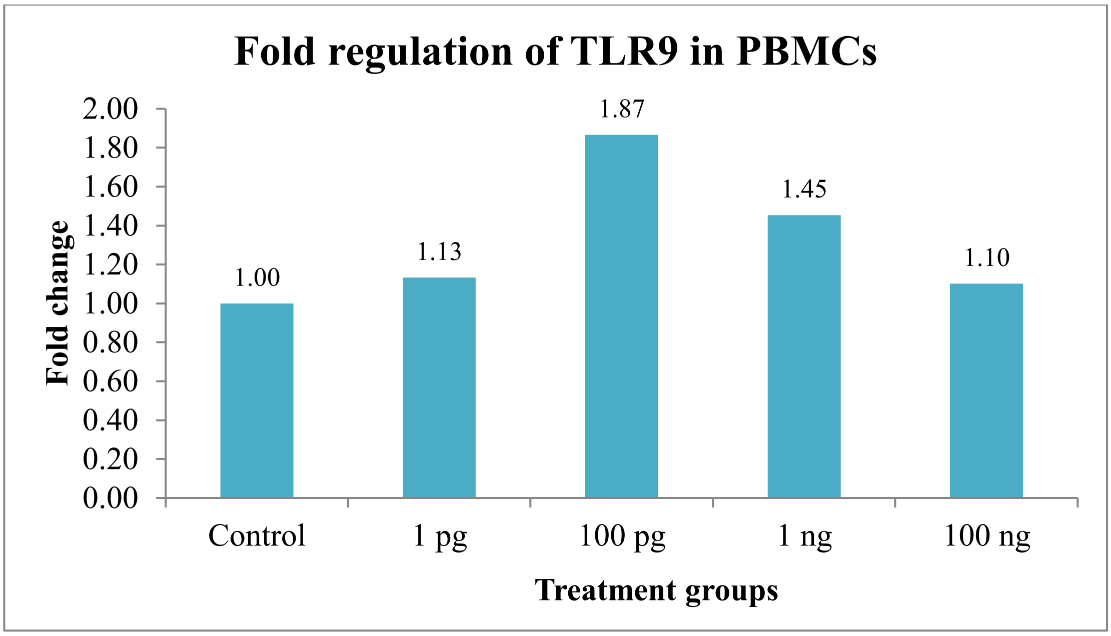


**Fig 25: Relative normalized expression of TLR 9 gene**

**Table 11: Relative normalized expression of TLR 10 gene**

| Treatment Group | TLR10 | GAPDH | ΔCq | ΔΔCq | Fold change 2^-ΔΔCq |
|---|---|---|---|---|---|
| Control | 27.11 | 20.66 | 6.45 | 0.00 | 1.00 |
| 1 pg | 26.78 | 20.86 | 5.92 | -0.53 | 1.44 |
| 100 pg | 26.76 | 21.47 | 5.29 | -1.16 | 2.23 |
| 1 ng | 26.91 | 21.09 | 5.82 | -0.63 | 1.55 |
| 100 ng | 27.00 | 21.03 | 5.97 | -0.48 | 1.39 |

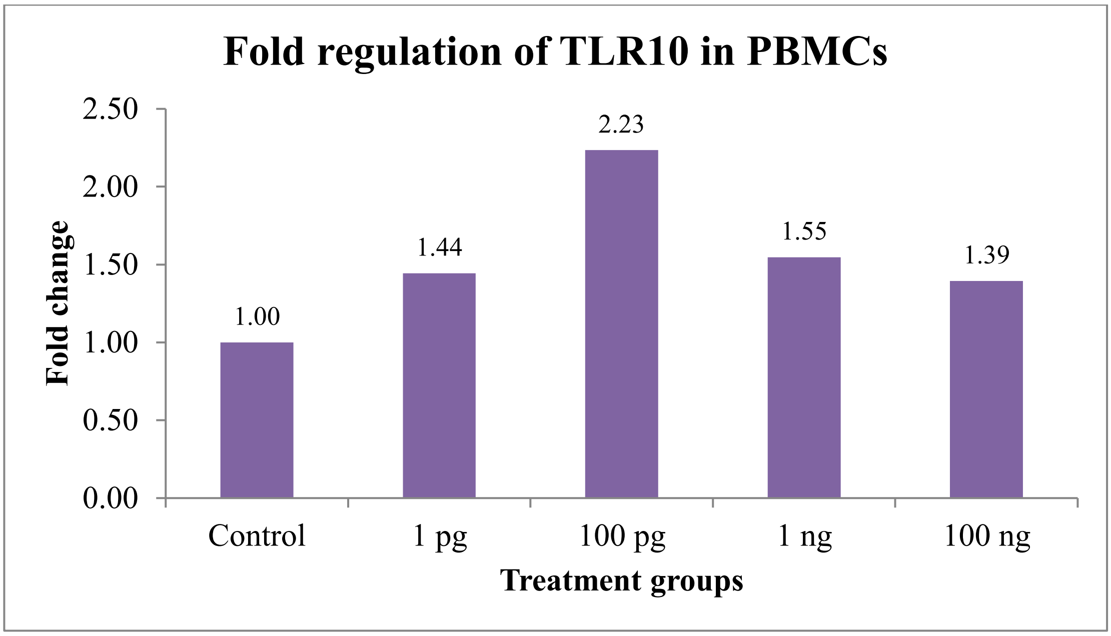


**Fig 26: Relative normalized expression of TLR 10 gene**

**Table 12: Relative normalized expression of MYD88 gene**

| Treatment Group | MYD88 | GAPDH | ΔCq | ΔΔCq | Fold change 2^-ΔΔCq |
|---|---|---|---|---|---|
| Control | 32.40 | 20.66 | 11.74 | 0.00 | 1.00 |
| 1 pg | 29.97 | 20.86 | 9.11 | -2.63 | 6.19 |
| 100 pg | 31.06 | 21.47 | 9.59 | -2.15 | 4.44 |
| 1 ng | 30.72 | 21.09 | 9.63 | -2.11 | 4.32 |
| 100 ng | 32.56 | 21.03 | 11.53 | -0.21 | 1.16 |

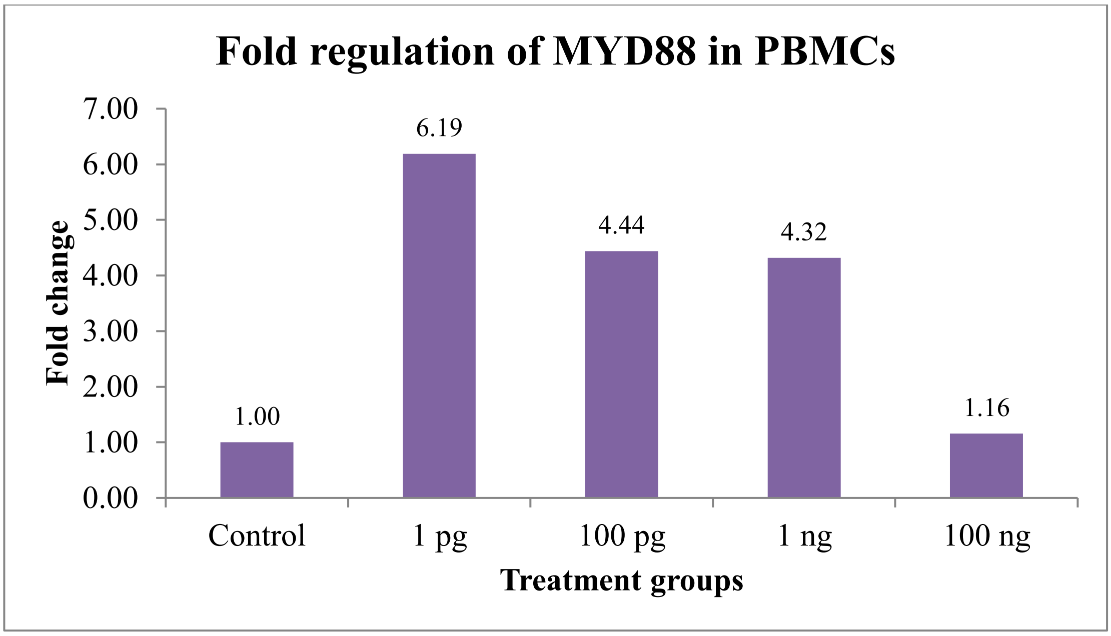


**Fig 27: Relative normalized expression of MYD88 gene**

**Table 13: Relative normalized expression of IRAK4 gene**

| Treatment Group | IRAK4 | GAPDH | ΔCq | ΔΔCq | Fold change 2^-ΔΔCq |
|---|---|---|---|---|---|
| Control | 22.57 | 20.66 | 1.91 | 0.00 | 1.00 |
| 1 pg | 23.94 | 20.86 | 3.08 | 1.17 | 0.44 |
| 100 pg | 25.05 | 21.47 | 3.58 | 1.67 | 0.31 |
| 1 ng | 23.52 | 21.09 | 2.43 | 0.52 | 0.70 |
| 100 ng | 23.76 | 21.03 | 2.73 | 0.82 | 0.57 |

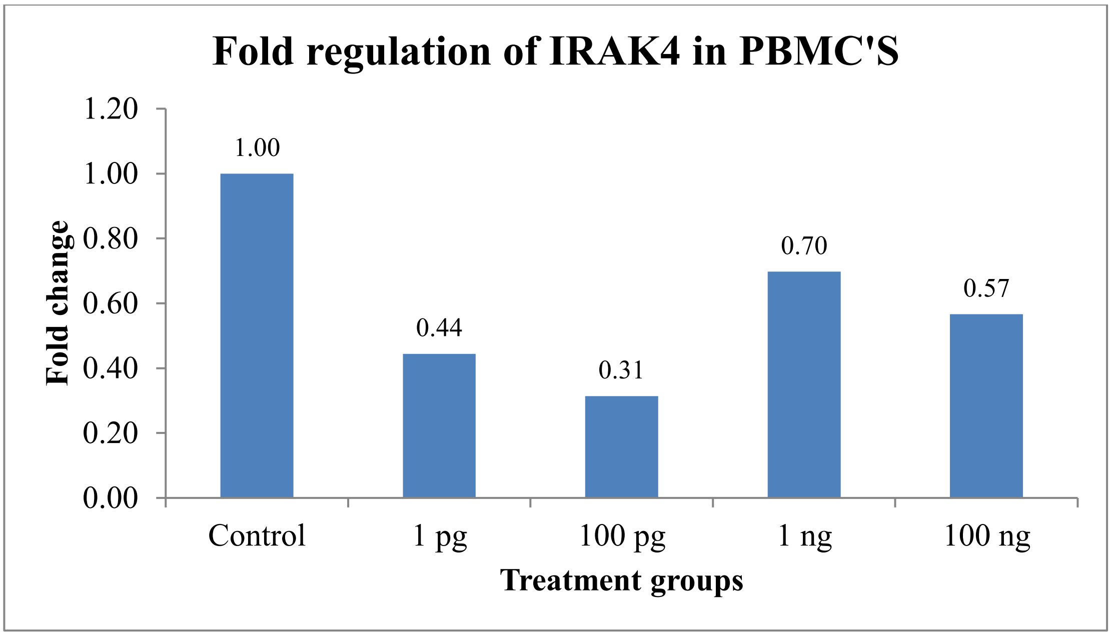


**Fig 28: Relative normalized expression of IRAK4 gene**

**Table 14: Relative normalized expression of TRAF3 gene**

| Treatment Group | TRAF3 | GAPDH | ΔCq | ΔΔCq | Fold change 2^-ΔΔCq |
|---|---|---|---|---|---|
| Control | 28.33 | 20.66 | 7.67 | 0.00 | 1.00 |
| 1 pg | 29.83 | 20.86 | 8.97 | 1.30 | 0.41 |
| 100 pg | 30.09 | 21.47 | 8.62 | 0.95 | 0.52 |
| 1 ng | 28.94 | 21.09 | 7.85 | 0.18 | 0.88 |
| 100 ng | 28.32 | 21.03 | 7.29 | -0.38 | 1.30 |

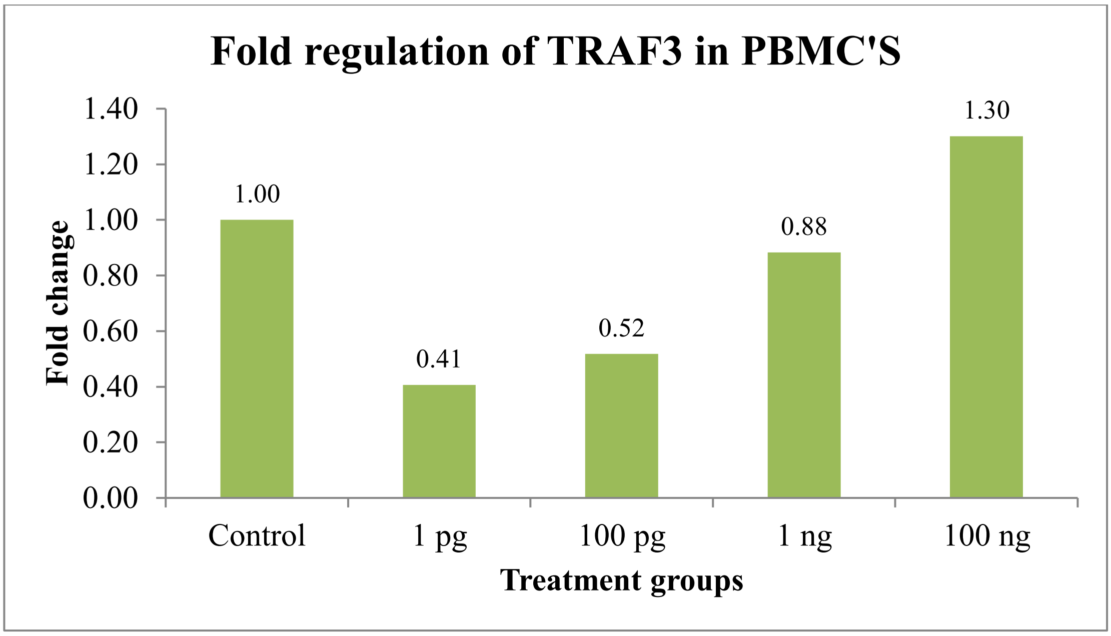


**Fig 29: Relative normalized expression of TRAF3 gene**

**Table 15: Relative normalized expression of TRAF6 gene**

| Treatment Group | TRAF6 | GAPDH | ΔCq | ΔΔCq | Fold change 2^-ΔΔCq |
|---|---|---|---|---|---|
| Control | 24.88 | 20.66 | 4.22 | 0.00 | 1.00 |
| 1 pg | 26.36 | 20.86 | 5.5 | 1.28 | 0.41 |
| 100 pg | 26.33 | 21.47 | 4.86 | 0.64 | 0.64 |
| 1 ng | 25.53 | 21.09 | 4.44 | 0.22 | 0.86 |
| 100 ng | 25.89 | 21.03 | 4.86 | 0.64 | 0.64 |

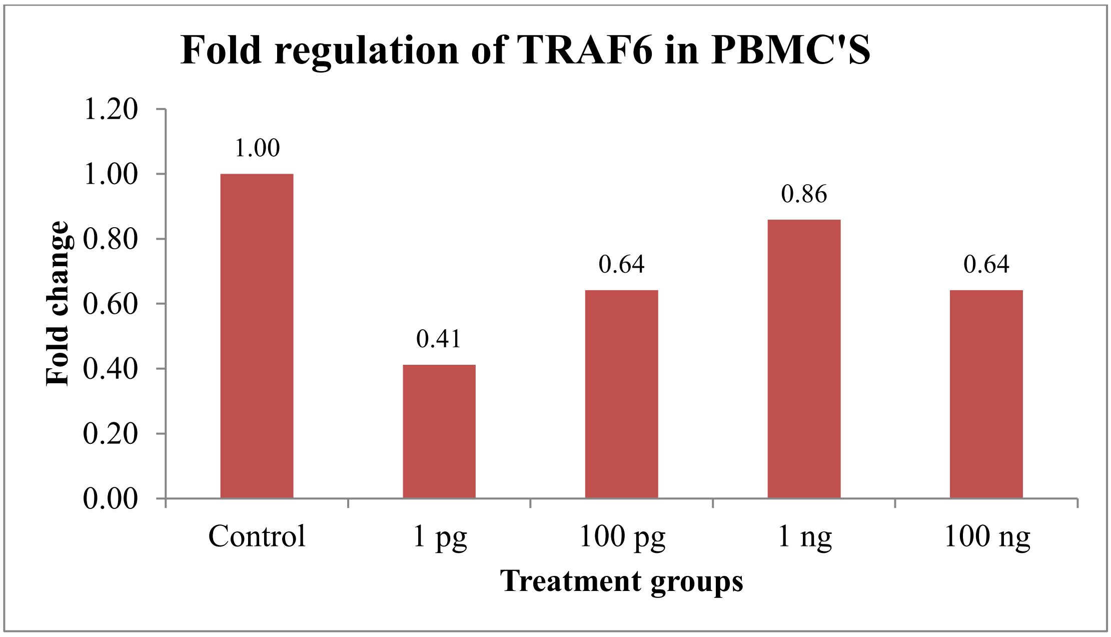


**Fig 30: Relative normalized expression of TRAF6 gene**

**Table 16: Relative normalized expression of TRIF gene**

| Treatment Group | TRIF | GAPDH | ΔCq | ΔΔCq | Fold change 2^-ΔΔCq |
|---|---|---|---|---|---|
| Control | 30.65 | 20.66 | 9.99 | 0.00 | 1.00 |
| 1 pg | 33.64 | 20.86 | 12.78 | 2.79 | 0.14 |
| 100 pg | 32.57 | 21.47 | 11.10 | 1.11 | 0.46 |
| 1 ng | 31.34 | 21.09 | 10.25 | 0.26 | 0.84 |
| 100 ng | 30.74 | 21.03 | 9.71 | -0.28 | 1.21 |

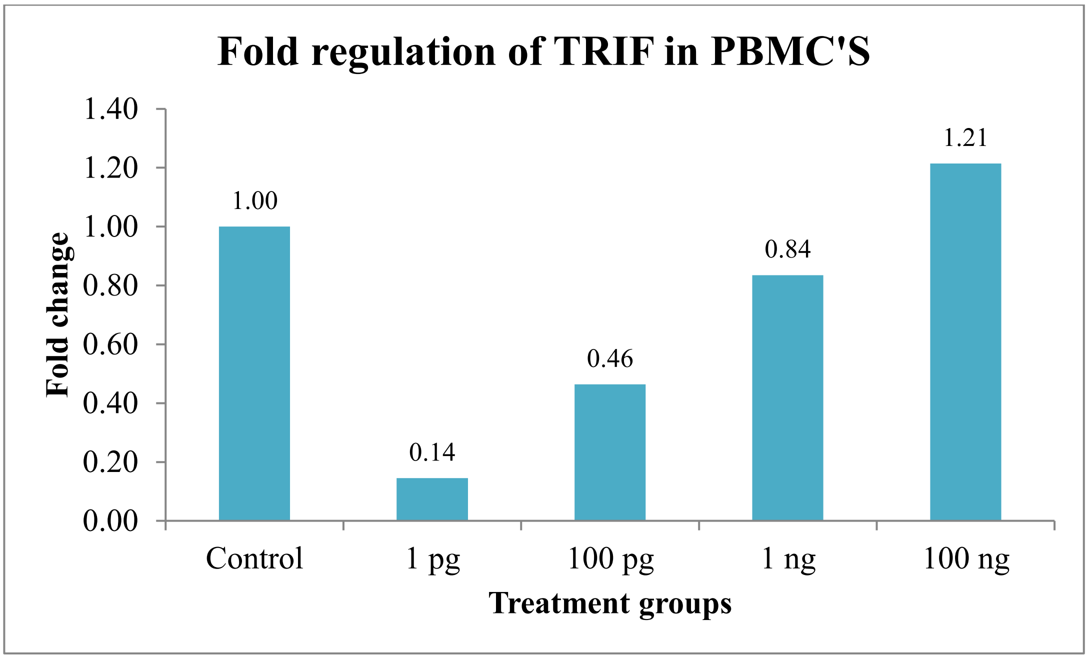


**Fig 31: Relative normalized expression of TRIF gene**

**Table 17. Summary of the results**

| Sr. No. | Treatment Details | Control | 1pg | 100pg | 1ng | 100ng |
|---|---|---|---|---|---|---|
| 1 | TLR1 | 1 | 1.99 | 2.81 | 2.31 | 1.91 |
| 2 | TLR2 | 1 | 1.72 | 2.38 | 2.06 | 1.68 |
| 3 | TLR3 | 1 | 1.53 | 1.53 | 1.57 | 1.01 |
| 4 | TLR4 | 1 | 0.67 | 0.97 | 0.95 | 0.85 |
| 5 | TLR5 | 1 | 5.54 | 2.91 | 2.83 | 1.1 |
| 6 | TLR6 | 1 | 4.5 | 3.43 | 2.66 | 2.14 |
| 7 | TLR7 | 1 | 1.64 | 2.69 | 2.07 | 1.69 |
| 8 | TLR8 | 1 | 2.43 | 6.54 | 1.88 | 1.33 |
| 9 | TLR9 | 1 | 1.13 | 1.87 | 1.45 | 1.1 |
| 10 | TLR10 | 1 | 1.44 | 2.23 | 1.55 | 1.39 |
| 11 | MYD88 | 1 | 6.19 | 4.44 | 4.32 | 1.16 |
| 12 | IRAK4 | 1 | 0.44 | 0.31 | 0.70 | 0.57 |
| 13 | TRAF6 | 1 | 0.41 | 0.64 | 0.86 | 0.64 |
| 14 | TRAF3 | 1 | 0.41 | 0.52 | 0.88 | 1.30 |
| 15 | TRIF | 1 | 0.14 | 0.46 | 0.84 | 1.21 |

**Comments**

The RT PCR analysis of target genes TLR1, TLR2, TLR3, TLR4, TLR5, TLR6, TLR7, TLR8, TLR9, and TLR10 were carried out and fold regulation was determined. The results suggested the significant up-regulation of TLR3, TLR6, TLR7, TLR8, TLR9, TLR10, MYD88, TRAF3, and TRIF with 1.01,2.14,1.69,1.33, 1.1 1.39, 1.16, 1.13, and 1.21 fold expressions respectively at the concentration of 100 ng when compared to control cells. Downregulation of IRAK4 and TRAF6, with 0.57 and 0.64 fold expression respectively at the concentration of 100 ng when compared to control cells.

---------------------------------------------------------------------------------------------
